# A next-generation, histological atlas of the human brain and its application to automated brain MRI segmentation

**DOI:** 10.1101/2024.02.05.579016

**Authors:** Adrià Casamitjana, Matteo Mancini, Eleanor Robinson, Loïc Peter, Roberto Annunziata, Juri Althonayan, Shauna Crampsie, Emily Blackburn, Benjamin Billot, Alessia Atzeni, Oula Puonti, Yaël Balbastre, Peter Schmidt, James Hughes, Jean C Augustinack, Brian L Edlow, Lilla Zöllei, David L Thomas, Dorit Kliemann, Martina Bocchetta, Catherine Strand, Janice L Holton, Zane Jaunmuktane, Juan Eugenio Iglesias

## Abstract

Magnetic resonance imaging (MRI) is the standard tool to image the human brain *in vivo*. In this domain, digital brain atlases are essential for subject-specific segmentation of anatomical regions of interest (ROIs) and spatial comparison of neuroanatomy from different subjects in a common coordinate frame. High-resolution, digital atlases derived from histology (e.g., Allen atlas [7], BigBrain [13], Julich [15]), are currently the state of the art and provide exquisite 3D cytoarchitectural maps, but lack probabilistic labels throughout the whole brain. Here we present *NextBrain*, a next - generation probabilistic atlas of human brain anatomy built from serial 3D histology and corresponding highly granular delineations of five whole brain hemispheres. We developed AI techniques to align and reconstruct ∼10,000 histological sections into coherent 3D volumes with joint geometric constraints (no overlap or gaps between sections), as well as to semi-automatically trace the boundaries of 333 distinct anatomical ROIs on all these sections. Comprehensive delineation on multiple cases enabled us to build *the first probabilistic histological atlas of the whole human brain*. Further, we created a companion Bayesian tool for automated segmentation of the 333 ROIs in any *in vivo* or *ex vivo* brain MRI scan using the *NextBrain* atlas. We showcase two applications of the atlas: automated segmentation of ultra-high-resolution *ex vivo* MRI and volumetric analysis of Alzheimer’s disease and healthy brain ageing based on ∼4,000 publicly available *in vivo* MRI scans. We publicly release: the raw and aligned data (including an online visualisation tool); the probabilistic atlas; the segmentation tool; and ground truth delineations for a 100 μm isotropic *ex vivo* hemisphere (that we use for quantitative evaluation of our segmentation method in this paper). By enabling researchers worldwide to analyse brain MRI scans at a superior level of granularity without manual effort or highly specific neuroanatomical knowledge, *NextBrain* holds promise to increase the specificity of MRI findings and ultimately accelerate our quest to understand the human brain in health and disease.

---

Magnetic resonance imaging (MRI) is arguably the most important tool to study the human brain *in vivo*. Its exquisite contrast between different types of soft tissue provides a window into the living brain without ionising radiation, making it suitable to healthy volunteers. Advances in magnet strength, data acquisition and image reconstruction methods [16-20] enable the acquisition of millimetre-resolution MRI scans of the *whole* brain in minutes. MRI can be acquired with different pulse sequences that image different tissue properties, including: neuroanatomy with structural acquisitions [21]; brain activity with functional MRI based on blood oxygenation [23]; vasculature with perfusion imaging and MR angiography [24-27]; or white matter fibres and microstructure with diffusion-weighted MRI [28,29].

Publicly available neuroimaging packages (FreeSurfer [30], FSL [31], SPM [32], or AFNI [33]) enable researchers to perform large-scale studies with thousands of scans [34-37] to study of healthy ageing, as well as a broad spectrum of brain diseases, such as Alzheimer’s, multiple sclerosis, or depression [38-41]. A core component of these neuroimaging packages is digital 3D brain atlases. These are reference 3D brain images that are representative of a certain population and can comprise image intensities, neuroanatomical labels, or both. We note that, due to its highly convoluted structure, the cerebral cortex is often modelled with specific atlases defined on surface coordinate systems [42,43] – rather than 3D images). We refer the reader to [44] for a comparative study.

Volumetric atlases are often computed by averaging data from a large cohort of subjects [45], but they may encompass as few as a single subject – particularly when built from labour-intensive modalities like histology [13]. Atlases enable aggregation of data from different subjects into a common coordinate frame (CCF), thus allowing analyses (e.g., group comparisons) as a function of spatial location. Atlases that include neuroanatomical labels also provide prior spatial information for analyses like automated image segmentation [46].

Most volumetric atlases, including those in neuroimaging packages, capitalise on the abundance of *in vivo* MRI scans acquired at ∼1 mm isotropic resolution. This voxel size is sufficient to represent information at the level of gyri, sulci, and subcortical nuclei. However, it is insufficient to study the brain at the subregion level, which is desirable as brain substructures (e.g., hippocampal subfields, nuclei of the thalamus) are known from animal models and *postmortem* human studies to have different function and connectivity [47]. This limitation can be circumvented with higher resolution images acquired *ex vivo*, typically with MRI or histology.

*Ex vivo* MRI has no motion artifacts and enables long acquisitions with voxels in the 100 μm range [3,48-50]. However, it fails to visualise cytoarchitecture and resolve many boundaries between brain areas. Histology, on the other hand, is a microscopic 2D modality that can visualise distinct aspects of cytoarchitecture using an array of stains – thus revealing neuroanatomy with much higher detail. Earlier versions of histological atlases were printed, often not digitised, and comprised only a small set of labelled sections. Representative examples include the Morel atlas of the thalamus and basal ganglia [51] or the Mai atlas of the whole brain [1] (Fig. 1A).

**Fig. 1:**
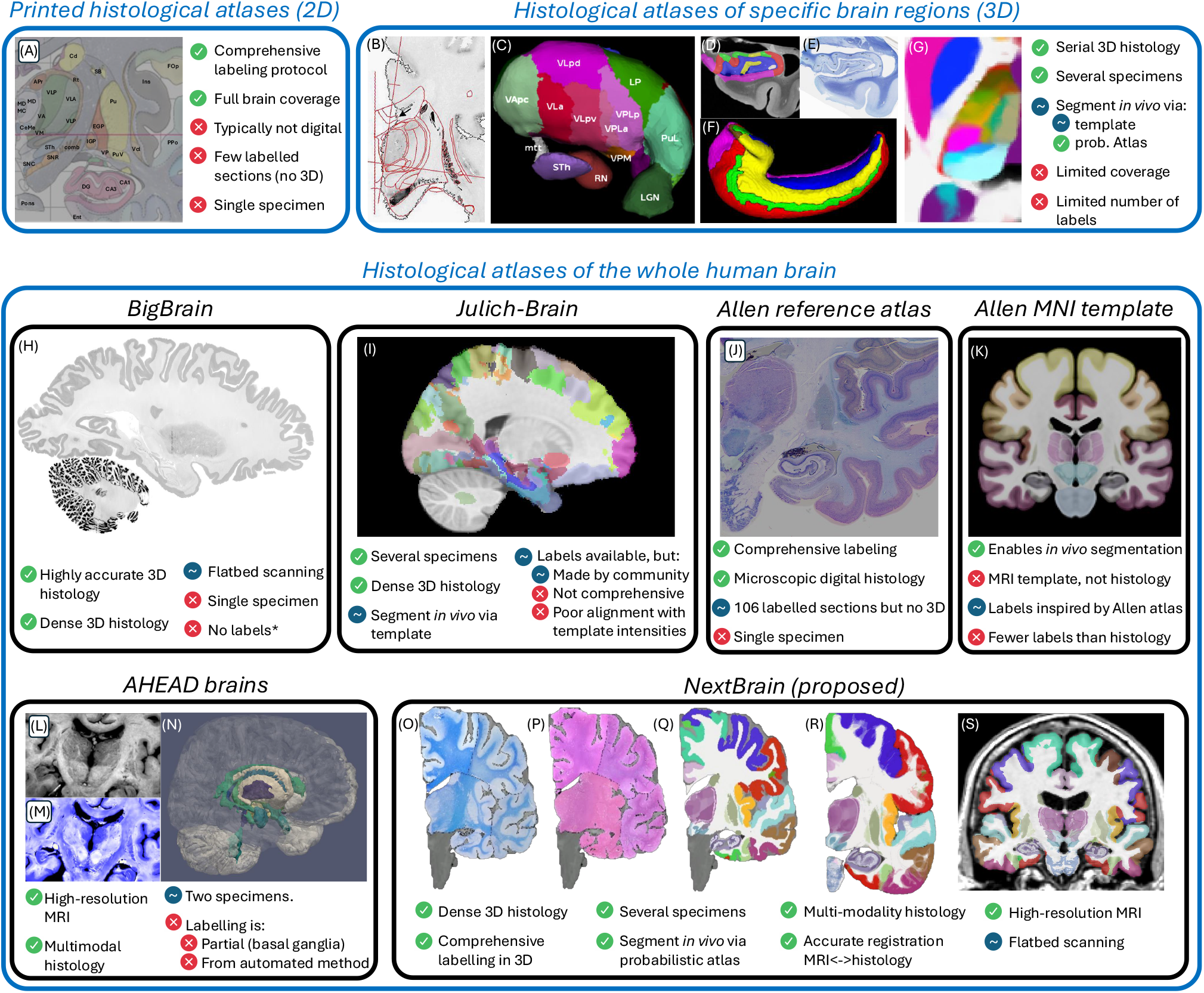
*NextBrain* in the context of histological atlases, with advantages (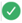), disadvantages (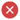), and neutral points. (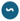). (A) Printed atlas [1] with a sparse set of manually traced sections [1]. (B-G) Histological atlases of specific ROIs with limited coverage: (B) Manually traced section of basal ganglia [8]; (C) 3D rendering of deterministic thalamic atlas [11]; (D-F) Traced MRI slice, histological section, and 3D rendering of hippocampal atlas [12]; and (G) Slice of our probabilistic atlas of the thalamus [14]. (H-N) Histological atlases of the whole human brain: (H) 3D reconstructed slice of BigBrain [13]; (I) Slice of Julich-Brain labels on MNI template; (J) Labelled histological section of the Allen reference brain [7]; (K) Labelling of MNI template with protocol inspired by (J); and (L-N) MRI, histology, and 3D rendering of AHEAD brains [22]. (O-S) Our new atlas *NextBrain* includes dense 3D histology (O-P) and comprehensive manual labels (Q) of five specimens, enabling the construction of a probabilistic atlas (R) that can be combined with Bayesian techniques to automatically label 333 ROIs in *in vivo* MRI scans (S).

While printed atlases are not useful for computational analysis, serial histology can be combined with image registration (alignment) methods to enable volumetric reconstruction of 3D histology [52], thus opening the door to creating 3D histological atlases. These have two major advantages over MRI atlases: *(i)* providing a more detailed CCF; and *(ii)* the ability to segment MRI scans at finer resolution – with potentially higher sensitivity and specificity to detect brain alterations caused by brain diseases or to measure treatment effects.

Earlier 3D histological atlases were limited in terms of anatomical coverage. Following the Morel atlas, two digital atlases of the basal ganglia and thalamus were presented [8,11] (Fig. 1B-C). To automatically obtain segmentations for living subjects, one needs to register their MRI scans with the histological atlases, which is difficult due to differences in image resolution and contrast between the two modalities. For this reason, the authors mapped the atlases to 3D MRI templates (e.g., the MNI atlas [53]) that can be more easily registered to *in vivo* images of other subjects. A similar atlas combining histological and MRI data was proposed for the hippocampus [12] (Fig. 1D-F). Our group presented a histological atlas of the thalamus [14] (Fig. 1G), but instead of using MNI as a stepping stone, we used Bayesian methods [54] to map our atlas to *in vivo* scans *directly*.

More recently, several efforts have aimed at the considerably bigger endeavour of building histological atlases of the whole human brain:

- BigBrain [13] comprises over 7,000 histological sections of a single brain, which were accurately reconstructed in 3D with an *ex vivo* MRI scan as reference (Fig. 1H). BigBrain paved the road for its follow-up Julich-Brain [15], which aggregates data from 23 individuals. A subset of 10 cases have been provided to the community for labelling, which has led to the annotation of 248 cytoarchitectonic areas as part of 41 projects. The maximum likelihood maps have been mapped to MNI space for *in vivo* MRI analysis [55], but have two caveats (Fig. 1I): they align poorly with the underlying MNI template, and subcortical annotations are only partial.
- The Allen reference brain [7] (Fig. 1J) has comprehensive anatomical annotations on high-resolution histology and is integrated with the Allen gene expression atlases. However, it only has delineations for a sparse set of histological sections of a single specimen (resembling a printed atlas). For 3D analysis of *in vivo* MRI, the authors have manually labelled the MNI template using a protocol inspired by their own atlas (Fig. 1K), but with a fraction of the labels and less accurately delineations – since they are made on MRI and not histology.
- The Ahead brains [22] (Fig. 1L-N) comprise quantitative MRI and registered 3D histology for two separate specimens. These have anatomical labels for a few dozen structures, but almost exclusively of the basal ganglia. Moreover, these labels were obtained from the MRI with automated methods, rather than manually traced on the high-resolution histology.

While these histological atlases of the whole brain provide exquisite 3D cytoarchitectural maps, interoperability with other datasets (e.g., gene expression), and some degree of MRI-histology integration, there are currently neither: *(i)* datasets with densely labelled 3D histology of the whole brain; nor *(ii)* probabilistic atlases built from such datasets, which would enable analyses such as Bayesian segmentation or CCF mapping of the whole brain at the subregion level.

In this article, we present *NextBrain*, a next-generation probabilistic atlas of the human brain built from comprehensively labelled, multi-modal 3D histology of five half brains (Fig. 1O-P). The full dataset comprises ∼10,000 sections stained with Hematoxylin and Eosin (H&E, which discerns cell nuclei vs cytoplasm) and Luxol Fast Blue (LFB, which enhances myelin). These sections were: *(i)* 3D-reconstructed with *ex vivo* MRI scans and highly customised image registration methods powered by artificial intelligence (AI); and *(ii)* densely segmented into 333 regions of interest (ROIs) with AI-enabled, semi-automated segmentation methods (Fig. 1Q). The 3D label maps are finally use to build a probabilistic atlas (Fig. 1R), which is combined with a Bayesian tool for automated segmentation of MRI scans (Fig. 1S).

As the first densely labelled probabilistic atlas of the human brain built from histology, *NextBrain* enables brain MRI analysis at a level of detail that was previously not possible. Our results showcase: the high accuracy of our 3D histology reconstructions; *NextBrain*’s ability to accurately segment MRI scans acquired *in vivo* or *ex vivo*; its ability to separate diseased and control subjects in an Alzheimer’s group study; and a volumetric study of healthy brain aging with unprecedented detail.

In addition to the atlas and companion segmentation tool, our public release of *NextBrain* includes: *(i)* The raw and registered images that were used to build the atlas, which are an invaluable resource for MRI signal modelling or histology registration studies; *(ii)* An online visualisation tool for these data, for educational and data inspection purposes; *(iii)* The source code and pipelines, which do not require any highly specialised equipment for intact coronally sliced full-brain procedures used at select sites like Allen or Julich (e.g., full-brain microtome, custom glass slides), thus enabling wide applicability; and *(iv)* our manual 3D segmentation of a publicly available 100 μm isotropic *ex vivo* scan [3] (used here for quantitative evaluation), which is a valuable resource in its own right, e.g., for ROI analysis in the *ex vivo* CCF of this scan, or for development and validation of segmentations methods.

## Densely labelled 3D histology of five human hemispheres

The *NextBrain* workflow is summarised in Fig. 2 and detailed in the Methods section. The first result of the pipeline (panels A-G) is a multimodal dataset with human hemispheres from five donors (three right, two left), including half cerebellum and brainstem. Each of the five cases comprises accurately aligned high-resolution *ex vivo* MRI, serial histology (H&E and LFB stains), and dense ground truth segmentations of 333 cortical and subcortical brain ROIs.

**Fig. 2:**
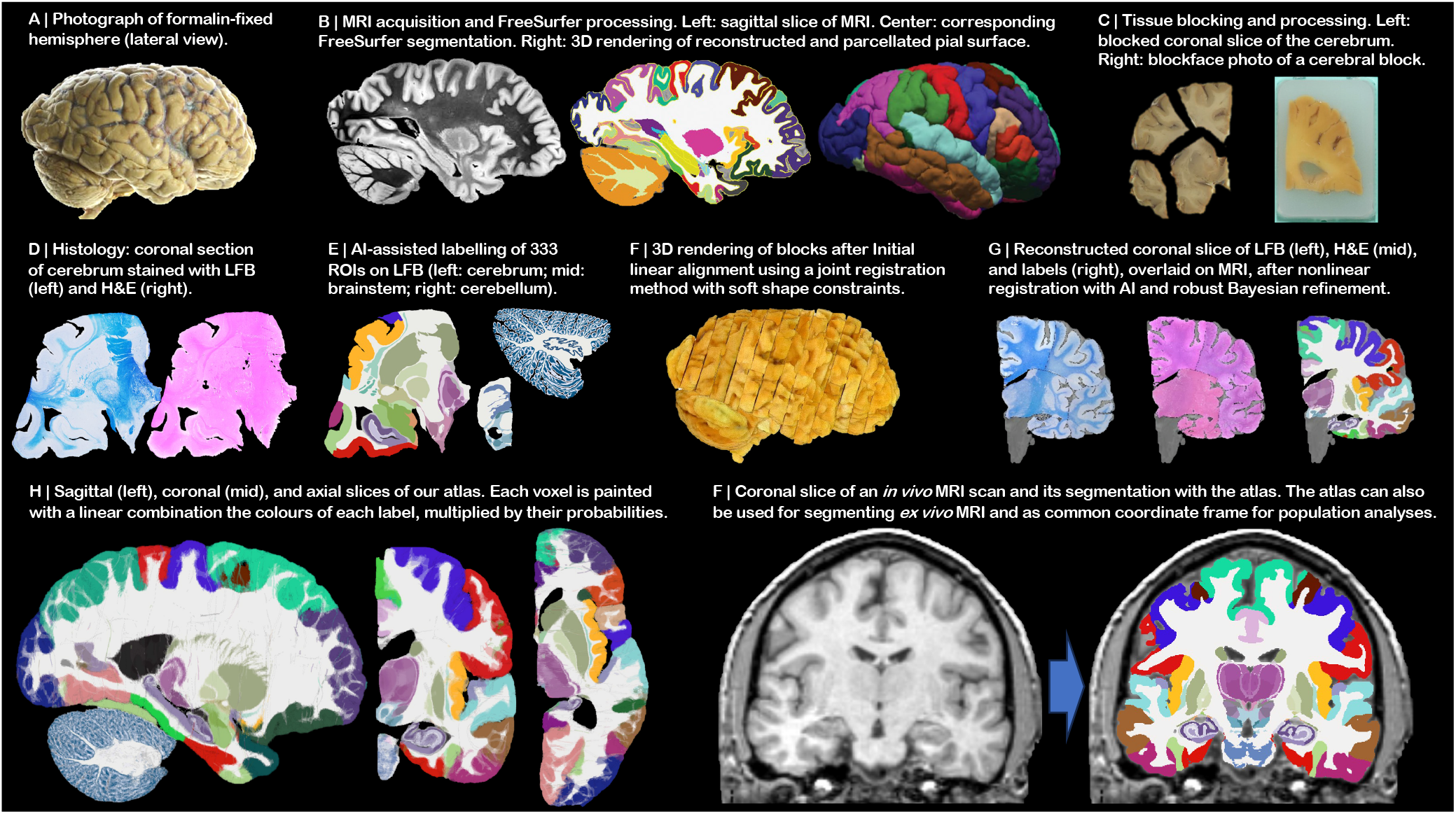
*NextBrain* workflow. (A) Photograph of formalin-fixed hemisphere. (B) High-resolution (400 μm) *ex vivo* MRI scan, FreeSurfer segmentation, and extracted pial surface (parcellated with FreeSurfer). (C) Tissue slabs and blocks, before and after paraffin embedding. (D) Section stained with H&E and LFB. (E) Semi-automated labelling of 333 ROIs on sections using an AI method [5]. (F) Initialization of affine alignment of tissue blocks using a custom registration algorithm that minimises overlap and gaps between blocks. (G) Refinement of registration with histology and nonlinear transform, using a combination of AI and Bayesian techniques [9,10]. (H) Orthogonal slices of 3D probabilistic atlas. (I) Automated Bayesian segmentation of an *in vivo* scan into 333 ROIs using the atlas.

Aligning the histology of a case is analogous to solving a 2,000-piece jigsaw puzzle in 3D, with the *ex vivo* MRI as reference (similar to the image on the box cover), and with pieces that are deformed by sectioning and mounting on glass slides – with occasional tissue folding or tearing. This problem falls out of the scope of existing inter-modality registration techniques [56], including slice-to-volume [57] and 3D histology reconstruction methods [52], which do not have to address the joint constraints of thousands of sections, acquired in nonparallel planes as part of different blocks.

Instead, we solve this challenging problem with a custom, state-of-the-art image registration framework (Fig. 3), which includes three components specifically developed for this project: *(i)* a differentiable regulariser that minimises overlap of different blocks and gaps in between [58]; *(ii)* an AI registration method that uses contrastive learning to provide highly accurate alignment of corresponding brain tissue across MRI and histology [10]; and *(iii)* a Bayesian refinement technique based on Lie algebra that guarantees the 3D smoothness of the reconstruction across modalities, even in the presence of outliers due to tissue folding and tearing [9]. We note that this is an evolution of our previously presented pipeline [6], which incorporates the aforementioned contrastive AI method and jointly optimises the affine and nonlinear transforms to achieve a 32% reduction in registration error (details below).

**Fig. 3:**
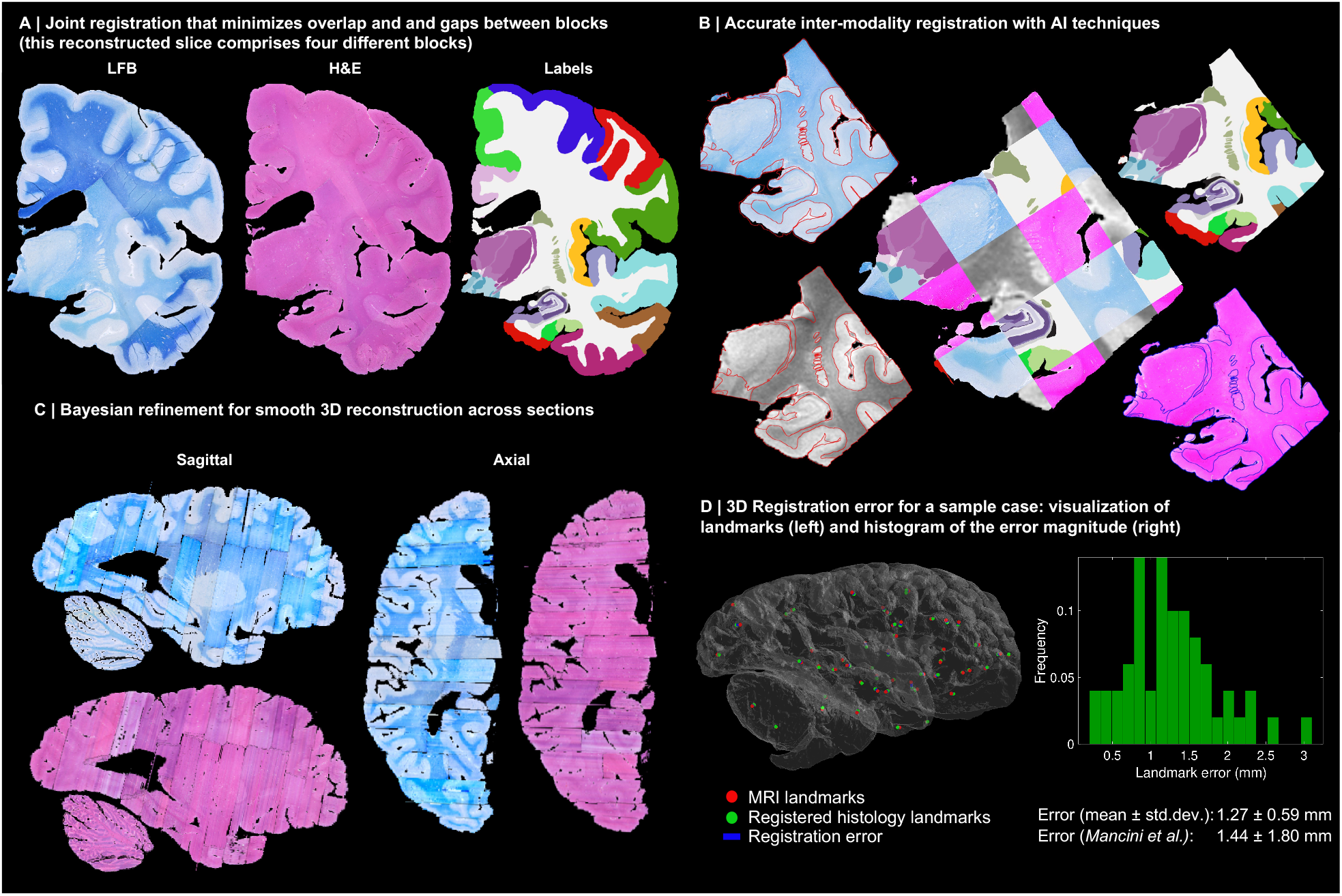
3D reconstruction of Case 1. (A) Coronal slice of 3D reconstruction; boundaries between blocks are noticeable from uneven staining. (B) Registered MRI, LFB, and H&E histology of a block, with tissue boundaries (traced on LFB) overlaid. (C) Orthogonal view of reconstruction, which is smooth thanks to the Bayesian refinement, and avoids gaps and overlaps thanks to the regulariser. (D) Visualization of 3D landmark registration error (left); histogram of its magnitude (right); and mean ± standard deviation (bottom), compared with our previous pipeline [6]. See Extended Data for results on the other cases. The average landmark error across all cases is 0.99mm (vs 1.45 for [6]).

Qualitatively, it is apparent from Fig. 3 that a very high level of accuracy is achieved for the spatial alignment, despite the non-parallel sections and distortions in the raw data. The regulariser effectively aligns the block boundaries in 3D without gaps or overlap (Fig. 3A-C), with minor discontinuities across blocks (e.g., in the temporal lobe). When the segmentations of different blocks are combined (Fig. 3A, right), the result is a smooth mosaic of ROI labels.

The AI-enabled registration across MRI and histological stains is exemplified in Fig. 3B. Overlaying the main ROI contours on the different modalities reveals the highly accurate alignment of the three modalities (MRI, H&E, LFB), even in convoluted regions of the cortex and the basal ganglia. The mosaic of modalities also highlights the accurate alignment at the substructural level, e.g., subregions of the hippocampus.

Fig. 3C shows the 3D reconstruction in orientations orthogonal to the main plane of sectioning (coronal). This illustrates not only the lack of gaps and overlaps between blocks, but also the smoothness that is achieved *within* blocks. This is thanks to the Bayesian refinement algorithm, which combines the best features of methods that: *(i)* align each section independently (high fidelity to the reference, but jagged reconstructions) and *(ii)* those that align sections to their neighbours (smooth reconstructions, but with “banana effect”, i.e., straightening of curved structures).

Quantitative results are presented in Fig. 3D, as well as in Extended Data Figs. 1-4D. The registration error, evaluated with 250 manually placed pairs of landmarks (known to be a better proxy for the registration error than similarity of label overlap metrics [59]), is 0.99 ± 0.51 mm – a considerable reduction with respect our previous pipeline [6], which yielded 1.45 ± 1.68 mm (Wilcoxon p=2×10_-22_). The spatial distribution of the error is further visualised with kernel regression in Extended Data Fig. 5, which shows that this distribution is fairly uniform, i.e., there is no obvious consistent pattern across cases.

Our pipeline is widely applicable as it produces accurate 3D reconstructions from blocked tissue in standard-sized cassettes, sectioned with a standard microtome. The computer code and aligned dataset is freely available in our public repository (see Data Availability). For educational and data inspection purposes, we have built an online visualisation tool for the multi-modality data, which is available at: github-pages.ucl.ac.uk/NextBrain.

Supplementary Video 1 illustrates the aligned data, which includes: *(i)* MRI at 400 μm isotropic resolution; *(ii)* aligned H&E and LFB histology digitised at 4 μm resolution (with 250 or 500 μm spacing, depending on the brain location); and *(iii)* ROI segmentations, obtained with a semi-automated AI method [5]. The ROIs comprise 34 cortical labels (following the Desikan-Killiany atlas [60]) and 299 subcortical labels (following different atlas for different brain regions; see the Methods section below and the supplement). This public dataset enables researchers worldwide to conduct their own studies not only in 3D histology reconstruction, but also other fields like: high-resolution segmentation of MRI or histology [61]; MRI-to-histology and histological stain-to-stain image translation [62]; deriving MRI signal models from histology [63]; and many others.

### A next-generation probabilistic atlas of the human brain

The labels from the five human hemispheres were coregistered and merged into a probabilistic atlas. This was achieved with a method that alternately registers the volumes to the estimate of the template, and updates the template via averaging [64]. The registration method is diffeomorphic [65] to ensure preservation of the neuroanatomic topology (e.g., ROIs do not split or disappear in the deformation process). Crucially, we use an initialization based on the MNI template, which serves two important purposes: preventing biases towards any of the cases (which would happen if we initialised with one of them); and “centring” our atlas on a well-established CCF computed from 305 subjects, which largely mitigates our relatively low number of cases. Since the MNI template is a greyscale volume, the first iteration of atlas building uses registrations computed with the *ex vivo* MRI scans. Subsequent iterations register labels directly with a metric based on the probability of the discrete labels according to the atlas [64].

Fig. 4 shows close-ups of orthogonal slices of the atlas, which models voxel-wide probabilities for the 333 ROIs on a 0.2mm isotropic grid. The resolution and detail of the atlas represents a substantial advance with respect to the SAMSEG atlas [2] currently in FreeSurfer (Fig. 4A). SAMSEG models 13 brain ROIs at 1 mm resolution and is, to the best of our knowledge, the most detailed probabilistic atlas that covers all brain regions. The figure also shows approximately corresponding slices of the manual labelling of the MNI atlas with the simplified Allen protocol [7]. Compared with *NextBrain*, this labelling is not probabilistic and does not include many histological boundaries that are invisible on the MNI template (e.g., hippocampal subregions, in violet). For this reason, it only has 138 ROIs – while *NextBrain* has 333.

**Fig. 4:**
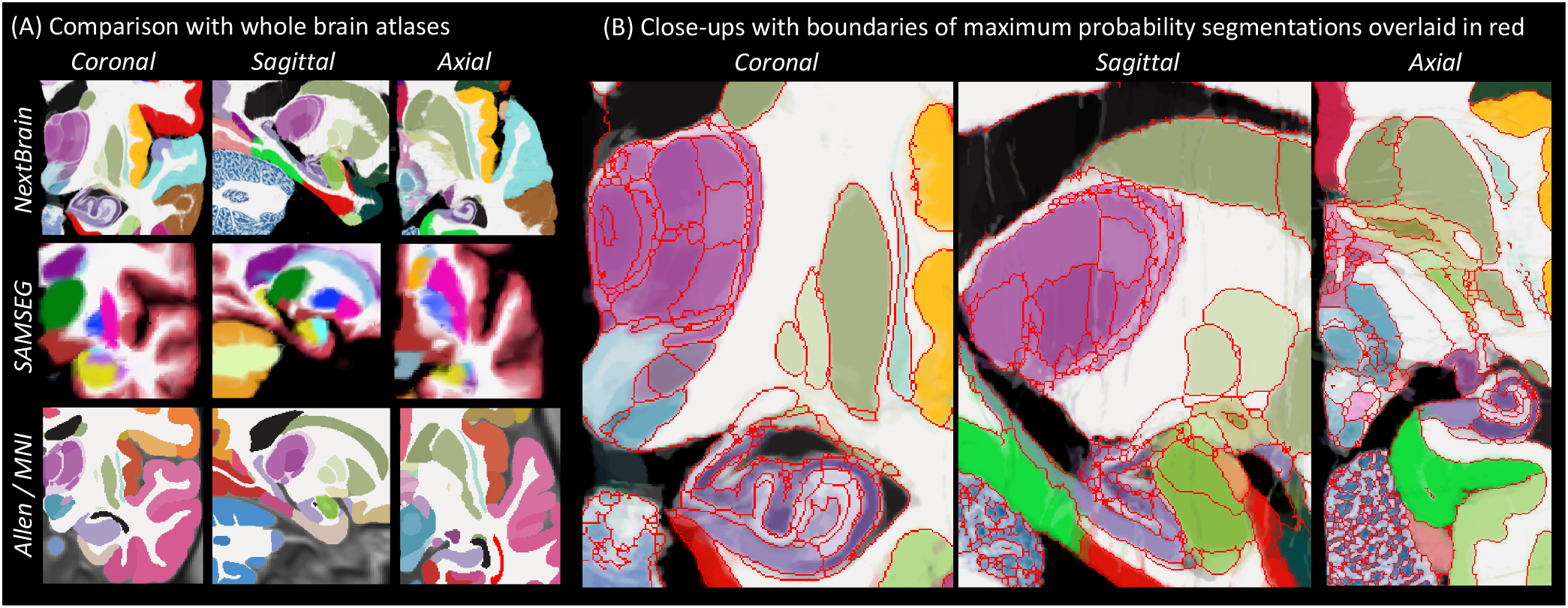
*NextBrain* probabilistic atlas. (A) Portions of the NextBrain probabilistic atlas (which has 333 ROIs), the SAMSEG atlas in FreeSurfer [2] (13 ROIs), and the manual labels of MNI based on the Allen atlas [7] (138 ROIs). (B) Close-up of three orthogonal slices of *NextBrain*. The colour coding follows the convention of the Allen atlas [7], where the hue indicates the structure (e.g., purple is thalamus, violet is hippocampus, green is amygdala) and the saturation is proportional to neuronal density. The colour of each voxel is a weighted sum of the colour corresponding to the ROIs, weighted by the corresponding probabilities at that voxel. The red lines separate ROIs based on the most probable label at each voxel, thus highlighting boundaries between ROIs of similar colour; we note that the jagged boundaries are a common discretization artefact of probabilistic atlases in regions where two or more labels mix continuously, e.g., the two layers of the cerebellar cortex.

A comprehensive comparison between and all digitised sections of the printed atlas by Mai & Paxinos [1] and approximately equivalent sections of the Allen reference brain and *NextBrain* is included in the supplement. The agreement between the three atlases is generally good, especially for the outer boundaries of the whole structures, e.g., the whole hippocampus, amygdala, or thalamus. Mild differences can be found in the delineation of sub-structures, both cortical and subcortical (e.g., subdivision of the accumbens), mainly due to: *(i)*the forced choice of applying arbitrary anatomical criteria in both atlases due to lack of contrast in smaller regions; *(ii)* different anatomical definitions; and *(iii)* the probabilistic nature of NextBrain. We emphasise that these differences are not exclusive to NextBrain, as they are also present between Mai-Paxinos and Allen.

Close-ups *NextBrain* slices centred on representative brain regions are shown in Fig. 4B, with boundaries between the ROIs (computed from the maximum likelihood segmentation) overlaid in red. These highlight the anatomical granularity of the new atlas, with dozens of subregions for areas such as the thalamus, hippocampus, amygdala, midbrain, etc. An overview of the complete atlas is shown in Supplementary Video 2, which illustrates the atlas construction procedure and flies through all the slices in axial, coronal, and sagittal view.

The probabilistic atlas is freely available as part of our segmentation module distributed with FreeSurfer. The maximum likelihood and colour-coded probabilistic maps (as in Fig. 4) can also be downloaded separately from our public repository, for quick inspection and educational purposes (see Data Availability). Developers of neuroimaging methods can freely capitalise on this resource, e.g., by extending the atlas via combination with other atlases or manually tracing new labels; or by designing their own segmentation methods using the atlas. Neuroimaging researchers can use the atlas for finegrained automated segmentation (as shown below), or as a highly detailed CCF for population analyses.

### Automated segmentation of ultra-high resolution *ex vivo* MRI

One of the new analyses that *NextBrain* enables is the automated fine-grained segmentation of ultra-high-resolution *ex vivo* MRI. Since motion is not a factor in *ex vivo* imaging, very long MRI scanning times can be used to acquire data at resolutions that are infeasible *in vivo*. One example is the publicly available 100 μm isotropic whole brain presented in [3], which was acquired in a 100-hour session on a 7T MRI scanner. Such datasets have huge potential in mesoscopic studies connecting microscopy with *in vivo* imaging [66].

Volumetric segmentation of ultra-high-resolution *ex vivo* MRI can be highly advantageous in neuroimaging in two different manners. First, by supplementing such scans (like the 100-micron brain) with neuroanatomical information that augments their value as atlases, e.g., as common coordinate frames or for segmentation purposes [67]. And second, by enabling analyses of *ex vivo* MRI datasets at scale (e.g., volumetry or shape analysis).

Dense manual segmentation of these datasets is practically infeasible, as it entails manually tracing ROIs on over 1,000 slices. Moreover, one typically seeks to label these images at a higher level of detail than *in vivo* (i.e., more ROIs of smaller sizes), which exacerbates the problem. One may utilise semi-automated methods like the AI-assisted technique we used in to build *NextBrain* (see previous section), which limits the manual segmentation to one every *N* slices [5] (*N=4* in this work). However, such a strategy only ameliorates the problem to a certain degree, as tedious manual segmentation is still required for a significant fraction of slices.

A more appealing alternative is thus automated segmentation. However, existing approaches have limitations, as they either: *(i)* were designed for 1 mm *in vivo* scans and do not capitalise on the increased resolution of *ex vivo* MRI [2,54]; or *(ii)* utilise neural networks trained with *ex vivo* scans but with a limited number of ROIs, due to the immense labelling effort that is required to generate the training data [61].

This limitation is circumvented by *NextBrain*: as a probabilistic atlas of neuroanatomy, it can be combined with well-established Bayesian segmentation methods (which are adaptive to MRI contrast) to segment ultrahigh-resolution *ex vivo* MRI scans into 333 ROIs. We have released in FreeSurfer an implementation that segments full brain scans in approximately 1h, using a desktop equipped with a graphics processing unit (GPU).

To quantitatively evaluate the segmentation method, we have created a ground truth segmentation of the public 100-micron brain [3], which we are publicly releasing as part of *NextBrain*. To make this burdensome task practical and feasible, we simplified it in five manners: *(i)* downsampling the data to 200 μm resolution; *(i)*labelling only one hemisphere; *(iii)* using the same semi-automated AI method as in *NextBrain* for faster segmentation; *(iv)* using FreeSurfer to automatically subdivide the cerebral cortex; and *(v)* labelling only a subset of 98 visible ROIs (see Supplementary Videos 3 and 4). Even with these simplifications, labelling the scan took over 100 hours of manual tracing effort.

We compared the ground truth labels with the automated segmentations produced by *NextBrain* using Dice overlap scores. Since the ground truth has fewer ROIs (particularly in the brainstem), we: *(i)* clustered the ROIs in the automated segmentation that correspond with the ROIs in the ground truth; and *(ii)* used a version of *NextBrain* in which the brainstem ROIs are simplified to better match those of the ground truth (with 264 labels instead of 333). The results are shown in Extended Data Table 1. As expected, there is a clear link between size and Dice. Larger ROIs like the cerebral white matter or cortex have Dice around 0.9. The smaller ROIs have lower Dice, but very few are below 0.4 – which is enough to *localize* ROIs. We note that the median Dice (0.667) is comparable with that reported by other Bayesian segmentation methods for brain subregions [68].

Sample slices and their corresponding automated and manual segmentations are shown in Fig. 5. The exquisite resolution and contrast of the dataset enables our atlas to accurately delineate a large number of ROIs with very different sizes, including small nuclei and subregions of the hippocampus, amygdala, thalamus, hypothalamus, midbrain, etc. Differences in label granularity aside, the consistency between the automated and ground truth segmentation is qualitatively very strong.

**Fig. 5:**
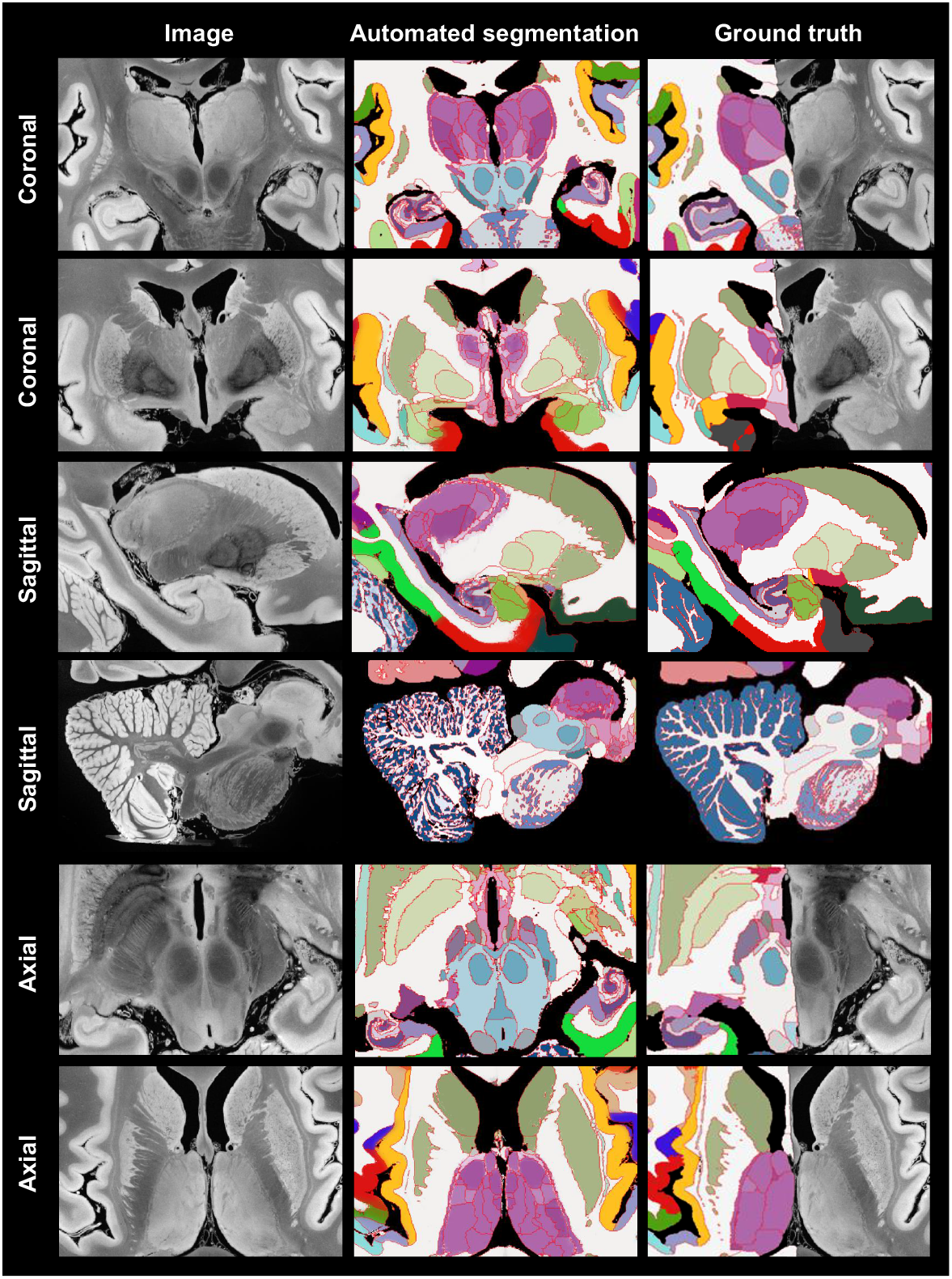
Automated Bayesian segmentation of publicly available ultra-high lution *ex vivo* brain MRI [3] using the simplified version of *NextBrain*, comparison with ground truth (only available for right hemisphere). We show two coronal, sagittal, and axial slices. The MRI was resampled to μm isotropic resolution for processing. As in previous figures, the segtation uses the Allen colour map [7] with boundaries overlaid in red. note that the manual segmentation uses a coarser labelling protocol.

To the best of our knowledge, this is the most comprehensive dense segmentation of a human brain MRI scan to date. As *ex vivo* datasets with tens of scans become available [61,69,70], our tool has great potential in augmenting mesoscopic studies of the human brain. Moreover, the labelled MRI that we are releasing has great potential in other neuroimaging studies, e.g., for training or evaluating segmentation algorithms; for ROI analysis in the high-resolution *ex vivo* space; or for volumetric analysis via registration-based segmentation.

### Fine-grained analysis of *in vivo* MRI

*NextBrain* can also be used to automatically segment *in vivo* MRI scans at the resolution of the atlas (200 μm isotropic), yielding an unprecedented level of detail. Scans used in research typically have isotropic resolution with voxel sizes ranging from 0.7 to 1.2mm, and therefore do not reveal all ROI boundaries with as much detail as ultra-high-resolution *ex vivo* MRI. However, many boundaries are still visible, including the external boundaries of brain structures (hippocampus, thalamus, etc.) and some internal boundaries, e.g., between the anteromedial and lateral posterior thalamus [14]. Bayesian segmentation capitalises on these visible boundaries and combines them with the prior knowledge encoded in the atlas to produce the full subdivision – albeit with lower reliability for the indistinct boundaries [49]. A sample segmentation is shown in Fig. 2F.

### Evaluation of segmentation accuracy

We first evaluated the *in vivo* segmentation quantitatively in two different experiments. First, we downsampled the *ex vivo* MRI scan from the previous section to 1 mm isotropic resolution (i.e., the standard resolution of *in vivo* scans); segmented it at 200 μm resolution; and computed Dice scores with the high-resolution ground truth. The results are displayed in Extended Data Table 1. The median Dice is 0.590, which is 0.077 lower than at 200 μm, but still fair for such small ROIs [68]. Moreover, most Dice scores remain 0.4, as for the ultra-high resolution, hinting that the priors can successfully provide a rough localization of internal boundaries, given the more visible external boundaries.

In a second experiment, we analysed the Dice scores produced by NextBrain in OpenBHB [4], a public metadataset with ∼1 mm isotropic T1-weighted scans of over 3,000 healthy individuals acquired at over 60 sites. Using FreeSurfer 7.0 as a silver standard, we computed Dice scores for our segmentations at the level of whole regions, i.e., the level of granularity provided by FreeSurfer. While these scores cannot assess segmentation accuracy at the subregion level, they do enable evaluation on a much larger multi-site cohort, as well as comparison with the Allen MNI template – the only competing histological (or rather, histology-inspired) atlas that can segment the whole brain *in vivo* (Fig. 1). The results (Extended Data Fig. 6) show that: *(i) NextBrain* consistently outperform the Allen MNI template, as expected from the fact that one atlas is probabilistic while the other is not; and *(ii) NextBrain* yields Dice scores in the range expected from Bayesian segmentation methods [2] – despite using only five cases, thanks to the excellent generalization ability of generative models [71].

### Application to Alzheimer’s disease (AD) classification

To further compare NextBrain with the Allen MNI template, we used an AD classification task based on linear discriminant analysis (LDA) of ROI volumes (corrected by age and intracranial volume). Using a simple linear classifier on a task where strong differences are expected allows us to use classification accuracy as a proxy for the quality of the input features, i.e., the ROI volumes derived from the automated segmentations. To enable direct comparison, we used a sample of 383 subjects from the ADNI dataset [72] (168 AD, 215 controls) that we used in previous publications [14,49,50].

Using the ROI volumes estimated by FreeSurfer 7.0 (which do *not* include subregions) yields and area under the receiver operating characteristic curve (AUROC) equal to 0.911, which classification accuracy of 85.4% at its elbow. The Allen MNI template exploits subregion information to achieve AUROC = 0.929 and 86.9% accuracy. The increased segmentation accuracy and granularity of NextBrain enables it to achieve AUROC = 0.953 and 90.3% accuracy – with a significant increase in AUROC with respect to the Allen MNI template (p = 0.01 for a DeLong test). This AUROC is also superior to those of specific *ex vivo* atlases we have presented in the prior work [14,49,50] – which range from 0.830 to 0.931

### Application to fine-grained signature of aging

We performed Bayesian segmentation with *NextBrain* on 705 subjects (aged 36-90, mean 59.6 years) from the Ageing HCP dataset [73], which comprises high-quality *in vivo* scans at 0.8mm resolution. We computed the volumes of the ROIs for every subject, corrected them for total intracranial volume (by division) and sex (by regression), and computed their Spearman correlation with age. We used the Spearman rather than Pearson correlation because, being rank-based, it is a better model for ageing trajectories as they are known to be nonlinear for wide age ranges [74,75].

The result of this analysis is, to the best of our knowledge, the most comprehensive map of regional ageing of the human brain to date (Fig. 6A and Extended Data Fig. 7A; see also full trajectories for select ROIs in Extended Data Fig. 8). Cortically, we found significant negative correlations with age in the prefrontal cortex (marked with ‘a’ on the figure) and insula (b), whilst the temporal (c) and parahippocampal cortices (d) did not yield significant correlation; this is consistent with findings from studies of cortical thickness [38,76]. The white matter (e) is known to decline steadily after ∼35 years [74,75], and such negative correlation is also detected by *NextBrain*. Other general ageing patterns at the whole structure level [74,75] are also successfully captured, such as a steady volume decrease of the caudate, thalamus, or putamen (f), or the volumetric reduction of the hippocampus, amygdala, or globus pallidus.

**Fig. 6:**
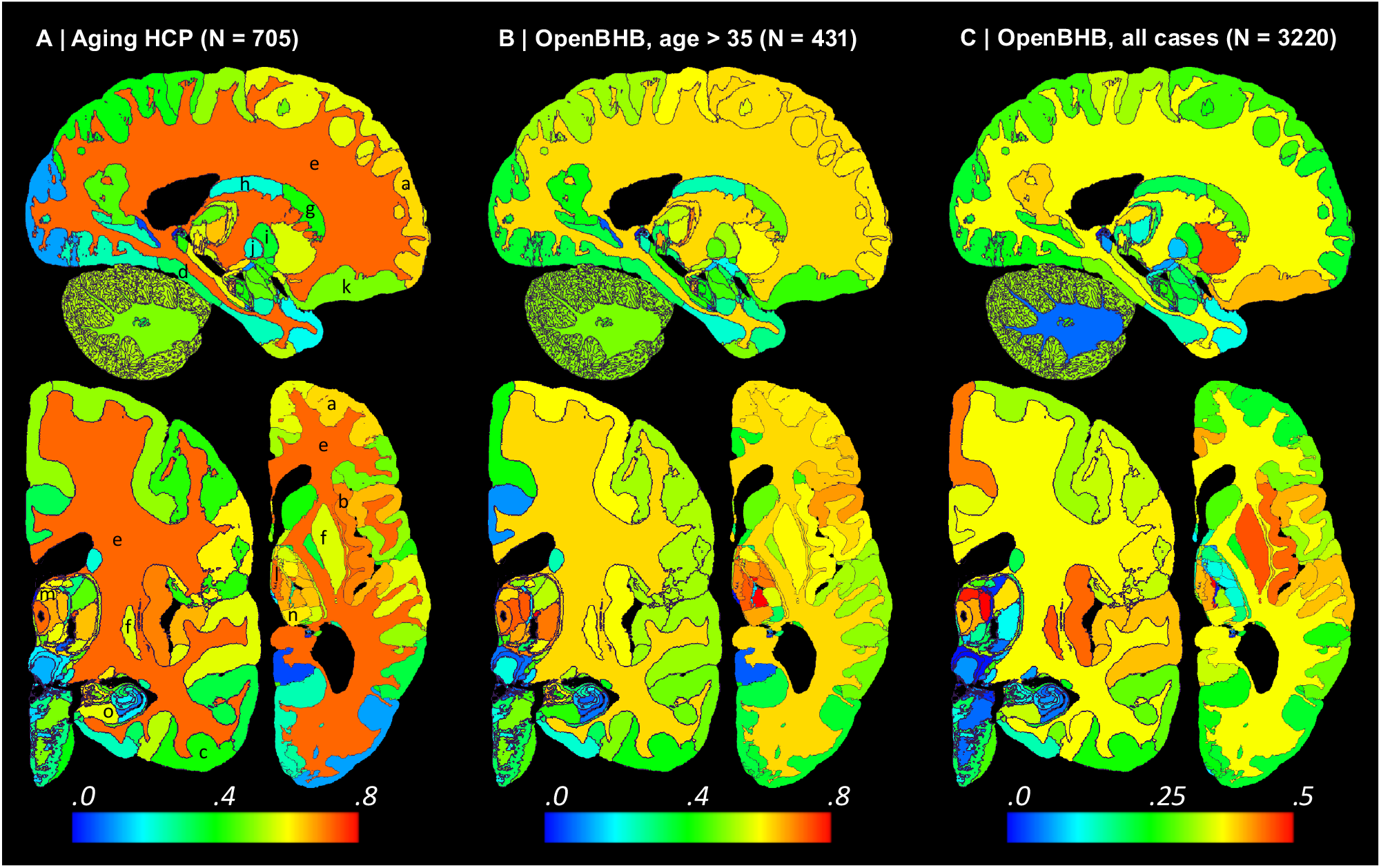
Absolute value of Spearman correlation for ROI volumes vs age derived from in vivo MRI scans: (A) Ageing HCP dataset (image resolution: .8mm isotropic; age range: 36-90 years; mean age: 59.6 years); please see main text for meaning of markers (letters). (B) OpenBHB dataset [4], restricted to subjects with ages over 35 years to match Ageing HCP (resolution 1 mm isotropic; age range: 36-86 years; mean age: 57.9 years). (C) Full OpenBHB dataset (age range: 6-86 years, mean age: 25.2 years); please note the different scale of the colour bar. The ROI volumes are corrected by intracranial volume (by division) and sex (by regression). Further slices are shown in Extended Data Fig. 6.

Importantly, *NextBrain* also unveils more granular patterns of the relationship between volumes and ageing within these regions. For example, the anterior caudate (g) showed a stronger negative correlation between age and volume than the posterior caudate (h). Similarly, the external segment of the globus pallidus (i) showed a stronger correlation than the internal segment (j) – an effect that was not observed in previous work studying the whole pallidum [77]. The ability to investigate separate subregions highlights a differential effect of ageing across brain networks, particularly a stronger effect on the regions of the limbic and prefrontal networks, given the correlations we found in the caudate head (g), insula (b), orbitofrontal cortex (k), amygdala, and thalamus [78]. Within the thalamus, the correlation is more significant in the mediodorsal (l), anteroventral (m), and pulvinar subnuclei (n), key regions in the limbic, lateral orbitofrontal and dorsolateral prefrontal circuits. In the hippocampus, subicular regions (o) correlate more strongly than the rest of the structure. The pattern of correlation strength is more homogeneous across subregions in the amygdala (key region in the limbic system), hypothalamus, and cerebellum.

We then revisited the OpenBHB dataset and performed the same regression analysis only for subjects older than 35 years, to match the age range of the Ageing HCP dataset (N=431, aged 36-86 years, mean 57.9 years). The results are shown in Fig. 6B and Extended Data Fig. 7B. Despite the differences in acquisition and the huge heterogeneity of the OpenBHB dataset, the results are highly consistent with those from HCP – but with slightly lower significance, possibly due to the increased voxel size (twice as big, since 1/0.8_3_ ≈ 2).

We also performed the same analysis with all 3,220 subjects in OpenBHB; see results in Fig. 6C and Extended Data Fig. 7C. For many regions, widening the age range to 6-86 years (mean age: 25.2) yields non-monotonic ageing curves and therefore weaker Spearman correlations. Therefore, these graphs highlight the regions whose volumes start decreasing with age the earliest, such as the putamen or medial thalamus. Many other patterns of association between age and ROI volumes remain very similar to those of the older populations (e.g., basal ganglia or hippocampus).

The segmentation code is publicly available in FreeSurfer: https://surfer.nmr.mgh.harvard.edu/fswiki/HistoAtlasSegmentation and can be run with a single line of code. This enables researchers worldwide to analyse their scans at a superior level of detail without manual effort or highly specific neuroanatomical knowledge.

## Discussion and Conclusion

*NextBrain* is a next-generation probabilistic human brain atlas, which is publicly available and distributed with a companion Bayesian segmentation tool and multi-modal dataset. The dataset itself is already a highly valuable resource: researchers have free access to both the raw and registered data, which they can use for their own research (e.g., in MRI signal modelling or registration), or to augment the atlas with new ROIs, e.g., by labelling them on the histology or MRI data and rebuilding the atlas. The atlas itself is a novel, high-resolution common coordinate frame for population analyses. The 3D segmentation of 100 μm *ex vivo* brain MRI scan [3] is a valuable complement to this (already very useful) resource. Finally, the Bayesian tool enables segmentation of *ex vivo* and *in vivo* MRI at an unprecedented level of granularity.

Due to its volumetric and semantic nature, NextBrain can be complemented by other segmentation methods and atlases that describe other aspects of the brain. For example, more accurate cortical segmentation and parcellation can be achieved with surface models [79]. We are currently working on models that combine neural networks with geometry processing to obtain laminar segmentations from both *in vivo* and *ex vivo* scans [80,81]. Surface placement will also warrant compatibility with cortical atlases obtained with multimodal data [43].

NextBrain is extensible: since all the data and code are publicly available, it is possible to download the data, modify (or extend) the manual annotations, and then rerun all the scripts to build a custom atlas. However, these tasks require domain expertise and compute power. Automatising this process to make it more accessible is desirable, but also quite a large engineering endeavour – and thus remains as future work.

The Bayesian segmentation tool in *NextBrain* is compatible with 1 mm isotropic scans, as illustrated by the Alzheimer’s and aging experiments. As with other probabilistic atlases, Bayesian segmentation can be augmented with models of pathology to automatically segment pathology, such as tumours [82] or white matter hyperintensities [83]. Importantly, *NextBrain*’s high level of detail enables us to fully take advantage of highresolution data, such as *ex vivo* MRI, ultra-high field MRI (e.g., 7T), and exciting new modalities like HiP-CT [84]. As high-quality 3D brain images become increasingly available, *NextBrain*’s ability to analyse them with superior granularity holds great promise to advance knowledge about the human brain in health and in disease.

## Methods

### Brain specimens

Hemispheres from five individuals (including half of the cerebrum, cerebellum, and brainstem), were used in this study, following informed consent to use the tissue for research and the ethical approval for research by the National Research Ethics Service (NRES) Committee London-Central. All hemispheres were fixed in 10% neutral buffered formalin (Fig. 2A). The laterality and demographics are summarised in Table 1 below; the donors were neurologically normal, but one case had an undiagnosed, asymptomatic tumour (diameter: ∼10mm) in the white matter, adjacent to the pars opercularis. This tumour did not pose issues in any of the processing steps described below.

**Table 1.** Demographics of the five cases used in this study.

| Case | Age at death | Sex | Laterality | Incidental findings |
| --- | --- | --- | --- | --- |
| Case 1 | 94 years | Female | Right | None |
| Case 2 | 84 years | Female | Right | None |
| Case 3 | 80 years | Male | Right | None |
| Case 4 | 79 years | Female | Left | Tumour adjacent to pars opercularis |
| Case 5 | 78 years | Male | Left | None |

### Data acquisition

Our data acquisition pipeline largely leverages our previous work [6]. We summarise it here for completeness; the reader is referred to the corresponding publication for further details.

#### MRI scanning

Prior to dissection, the hemispheres were scanned on a 3T Siemens MAGNETOM Prisma scanner. The specimens were placed in a container filled with Fluorinert (perfluorocarbon), a proton-free fluid with no MRI signal that yields excellent *ex vivo* MRI contrast and does not affect downstream histological analysis [85]. The MRI scans were acquired with a T2weighted sequence (optimised long echo train 3D fast spin echo [86]) with parameters: TR = 500 ms, TEeff = 69ms, BW = 558 Hz/Px, echo spacing = 4.96ms, echo train length = 58, 10 averages, with 400 μm isotropic resolution, acquisition time for each average = 547s, total scanning time = 91 min. These scans were processed with a combination of SAMSEG [2] and the FreeSurfer 7.0 cortical stream [79] to bias field correct the images, generate rough subcortical segmentations, and obtain white matter and pial surfaces with corresponding parcellations according to the Desikan-Killiany atlas [60] (Fig. 2B).

#### Dissection

After MRI scanning, each hemisphere is dissected to fit into standard 74×52mm cassettes. First, each hemisphere was split into cerebrum, cerebellum, and brainstem. Using a metal frame as a guide, these were subsequently cut into 10mm-thick slices in coronal, sagittal, and axial orientation, respectively. These slices were photographed inside a rectangular frame of known dimensions for pixel size and perspective correction; we refer to these images as “whole slice photographs.” While the brainstem and cerebellum slices all fit into the cassettes, the cerebrum slices were further cut into as many blocks as needed. “Blocked slice photographs” were also taken for these blocks (Fig. 2C, left).

#### Tissue processing and sectioning

After standard tissue processing steps, each tissue block was embedded in paraffin wax and sectioned with a sledge microtome at 25*μ*m thickness. Before each cut, a photograph was taken with a 24MPx Nikon D5100 camera (ISO = 100, aperture = f/20, shutter speed = automatic) mounted right above the microtome, pointed perpendicularly to the sectioning plane. These photographs (henceforth “blockface photographs”) were corrected for pixel size and perspective using fiducial markers. The blockface photographs have poor contrast between grey and white matter (Fig. 2C, right) but also negligible nonlinear geometric distortion, so they can be readily stacked into 3D volumes. A 2D convolutional neural network (CNN) pretrained on the ImageNet dataset [87] and fine-tuned on 50 manually labelled examples was used to automatically produce binary tissue masks for the blockface images.

#### Staining and digitisation

We mounted on glass slides and stained two consecutive sections every *N* (see below), one with Hematoxylin and Eosin (H&E) and one with Luxol Fast Blue (LFB); see Fig. 2D. The sampling interval was *N=10* (i.e., 250*μ*m) for blocks that included subcortical structures in the cerebrum, medial structures of the cerebellum, or brainstem structures. The interval was *N=20* (500*μ*m) for all other blocks. All stained sections were digitised with a flatbed scanner at 6,400 DPI resolution (pixel size: 3.97*μ*m). Tissue masks were generated using a 2D CNN similar to the one used for blockface photographs (pretrained on ImageNet and finetuned on 100 manually labelled examples).

### Dense labelling of histology

Segmentations of 333 ROIs (34 cortical, 299 subcortical) were made by authors ER, JA, and EB (with guidance from DK, MB, JZ, and JCA) for all the LFB sections, using a combination of manual and automated techniques (Fig. 2E). The general procedure to label each block was: *(i)* produce an accurate segmentation for one of every four sections; *(ii)* run SmartInterpol [5] to automatically segment the sections in between; and *(iii)* manually correct these automatically segmented sections when needed. SmartInterpol is a dedicated AI technique that we have developed specifically to speed up segmentation of histological stacks in this project.

To obtain accurate segmentations on sparse sections, we used two different strategies depending on the brain region. For the blocks containing subcortical or brainstem structures, ROIs were manually traced from scratch using a combination of ITK-SNAP [88] and FreeSurfer’s viewer “Freeview”. For cerebellum blocks, we first trained a 2D CNN (a U-Net [89]) on 20 sections on which we had manually labelled the white matter and the molecular and granular layers of the cortex. The CNN was then run on the (sparse) sections, and the outputs manually corrected. This procedure saves a substantial amount of time, since manually tracing the convoluted shape of the arbor vitae is extremely time consuming. For the cortical cerebrum blocks, we used a similar strategy as for the cerebellum, labelling the tissue as either white or grey matter. The subdivision of the cortical grey matter into parcels was achieved by taking the nearest neighbouring cortical label from the aligned MRI scan (details on the alignment below).

The manual labelling followed neuroanatomical protocols based on different brain atlases, depending on the brain region. Further details on the specific delineation protocols are provided in the Supplementary Methods. The general ontology of the 333 ROIs is based on the Allen reference brain [7], and is provide in a spreadsheet as part of the Supplementary Data.

### 3D histology reconstruction

3D histology reconstruction is the inverse problem of reversing all the distortion that brain tissue undergoes during acquisition, in order to reassemble a 3D shape that accurately follows the original anatomy. For this purpose, we used a framework with four modules.

#### Initial blockface alignment

In order to roughly initialise the 3D reconstruction, we relied on the stacks of blockface photographs. Specifically, we used our previously presented hierarchical joint registration framework [58] that seeks to: *(i)* align each block to the MRI with a similarity transform, by maximising the normalised crosscorrelation of their intensities; while *(ii)* discouraging overlap between blocks or gaps in between, via a differentiable regulariser. The similarity transforms allowed for rigid deformation (rotation, translation), as well as isotropic scaling to model the shrinking due to tissue processing. The registration algorithm was initialised with transforms derived from the whole slice, blocked slice, and blockface photographs (see details in [6]). The registration was hierarchical in the sense that groups of transforms were forced to share the same parameters in the earlier iterations of the optimisation, to reflect our knowledge of the cutting procedure. In the first iterations, we clustered the blocks into three groups: cerebrum, cerebellum, and brainstem. In the following iterations, we clustered the cerebral blocks that were cut from the same slice, and allowed translations in all directions, in-plane rotation, and global scaling. In the final iterations, each block alignment was optimised independently. The numerical optimisation used the LBFGS algorithm [90]. The approximate average error after this procedure was ∼2mm [58]. A sample 3D reconstruction is shown in Fig. 2F.

#### Refined alignment with preliminary nonlinear model

Once a good initial alignment is available, we can use the LFB sections to refine the registration. These LFB images have exquisite contrast (Fig. 2D) but suffer from nonlinear distortion – rendering the good initialization from the blockface images crucial. The registration procedure was nearly identical to that of the blockface, with two main differences. First, the similarity term used the local (rather than global) normalised cross-correlation function [91] in order to handle uneven staining across sections. Second, the deformation model and optimisation hierarchy were slightly different since nonlinear registration benefits from more robust methods. Specifically: The first two levels of optimisation were the same, with blocks grouped into cerebrum/cerebellum/brainstem (first level) or cerebral slices (second level), and optimisation of similarity transforms. The third level (i.e., each block independently) was subdivided into four stages in which we optimised transforms with increasing complexity, such that the solution of every level of complexity serves as initialisation to the next. In the first and simplest level, we allowed for translations in all directions, in-plane rotation and global scaling (5 parameters per block). In the second level, we added a different scaling parameter in the normal direction of the block (6 parameters per block). In the third level, we allowed for rotation in all directions (8 parameters per block). In the fourth and final level, we added to every section in every block a nonlinear field modelled with a grid of control points (10mm spacing) and interpolating B-splines. This final deformation model has approximately 100,000 parameters per case (∼100 parameters per section, times ∼1,000 LFB sections).

#### Nonlinear AI registration

We seek to produce final nonlinear registrations that are accurate, consistent with each other, and robust against tears and folds in the sections. We capitalise on Synth-by-Reg (SbR [10]), an AI tool for multimodal registration that we have recently developed, to register histological sections to MRI slices resampled to the plane of the histology (as estimated by the linear alignment). SbR exploits the facts that: *(i)* intra-modality registration is more accurate than inter-modality registration [92]; and *(ii)* there is a correspondence between histological sections and MRI slices, i.e., they represent the same anatomy. In short, SbR trains a CNN to make histological sections look like MRI slices (a task known as style transfer [93]), using a second CNN that has been previously trained to register MRI slices to each other. The style transfer relies on the fact that only good MRI synthesis will yield a good match when used as input to the second CNN, combined with a contrastive loss [94] that prevents blurring and content shift due to overfitting. SbR produces highly accurate deformations parameterised as stationary velocity fields (SVF [95]).

#### Bayesian refinement

Running SbR for each stain and section independently (i.e., LFB to resampled MRI, and H&E to resampled MRI) yields a reconstruction that is jagged and sensitive to folds and tears. One alternative is to register each histological section to each neighbour directly, which achieves smooth reconstructions but incurs the so-called “banana effect”, i.e., a straightening of curved structures [52]. We have proposed a Bayesian method that yields smooth reconstructions without banana effect [9]. This method follows an overconstrained strategy, by computing registrations between: LFB and MRI; H&E and MRI; H&E and LFB; each LFB section and the two nearest neighbours in either direction across the stack; each H&E section and its neighbours; and each MRI slice and its neighbours. For a stack with *S* sections, this procedure yields 15*xS-18* registrations, while the underlying dimensionality of the spanning tree connecting all the images is just 3xS-1. We use a probabilistic model of SVFs to infer the most likely spanning tree given the computed registrations, which are seen as noisy measurements of combinations of transforms in the spanning tree. The probabilistic model uses a Laplace distribution, which relies on L1 norms and is thus robust to outliers. Moreover, the properties of SVFs enable us to write the optimization problem as a linear program, which we solve with a standard simplex algorithm [96]. The result of this procedure was a 3D reconstruction that is accurate (it is informed by many registrations), robust, and smooth (Figures 2G and 3).

### Atlas construction

The transforms for the LFB sections produced by the 3D reconstructions were applied to the segmentations to bring them into 3D space. Despite the regulariser from [58], minor overlaps and gaps between blocks still occur. The former were resolved by selecting the label which is furthest inside the corresponding ROI. For the latter, we used our previously developed smoothing approach [14].

Given the low number of available cases, we combined the left (2) and right (3) hemispheres into a single atlas. This was achieved by flipping the right hemispheres and computing a probabilistic atlas of the left hemisphere using an iterative technique [64]. To initialise the procedure, we registered the MRI scans to the MNI atlas [53] with the right hemisphere masked out, and averaged the deformed segmentations to obtain an initial estimate of the probabilistic atlas. This first registration was based on intensities, using a local normalised cross-correlation loss. From that point on, the algorithm operates exclusively on the segmentations.

Every iteration of the atlas construction process comprises two steps. First, the current estimate of the atlas and the segmentations are co-registered one at the time, using: *(i)* a diffeomorphic deformation model based on SVFs parameterised by grids of control points and B-splines (as implemented in NiftyReg [97]), which preserves the topology of the segmentations; *(ii)* a data term, which is the log-likelihood of the label at each voxel according to the probabilities given by the deformed atlas (with a weak Dirichlet prior to prevent logs of zero); and *(iii)* a regulariser based on the bending energy of the field, which encourages regularity in the deformations. The second step of each iteration updates the atlas by averaging the segmentations. The procedure converged (negligible change in the atlas) after five iterations. Slices of the atlas are shown in Figs. 2H and 4.

### Bayesian segmentation

Our Bayesian segmentation algorithm builds on well-established methods in the neuroimaging literature [54,98,99]. In short, the algorithm jointly estimates a set of parameters that best explain the observed image in light of the probabilistic atlas, according to a generative model based on a Gaussian mixture model (GMM) conditioned on the segmentation, combined with a model of bias field. The parameters include the deformation of the probabilistic atlas, a set of coefficients describing the bias field, and the means, variances, and weights of the GMM. The atlas deformation is regularised in the same way as the atlas construction (bending energy, in our case) and is estimated via numerical optimisation with LBFGS. The bias field and GMM parameters are estimated with the Expectation Maximisation algorithm [100].

Compared with classical Bayesian segmentation methods operating at 1 mm resolution with just a few classes (e.g., SAMSEG [2], SPM [54]), our proposed method has several distinct features:

- Since the atlas only describes the left hemisphere, we use a fast deep learning registration method (EasyReg [101]) to register the input scan to MNI space, and use the resulting deformation to split the brain into two hemispheres that are processed independently.
- Since the atlas only models brain tissue, we run SynthSeg [102] on the input scan to mask out the extracerebral tissue.
- Clustering ROIs into tissue types (rather than letting each ROI have its own Gaussian) is particularly important, given the large number of ROIs (333). The user can specify the clustering via a configuration file; by default, our public implementation uses a configuration with 15 tissue types, tailored to *in vivo* MRI segmentation.
- The framework is implemented using the PyTorch package, which enables it to run on GPUs and curbs segmentation run times to about half an hour per hemisphere.

Sample segmentations with this method can be found in Figures 2H (*in vivo*) and 5 (*ex vivo*).

### Labelling of ultra-high resolution *ex vivo* brain MRI and simplified version of *NextBrain* atlas

In order to quantitatively assess the accuracy of our segmentation method on the ultra-high resolution *ex vivo* scan, we produced a gold standard segmentation of the publicly available 100 μm scan [3] as follows. First, we downsampled the data to 200 μm resolution and discarded the left hemisphere, to alleviate the manual labelling requirements. Next, we used *Freeview* to manually label from scratch one coronal slice every 10; we labelled as many regions from the histological protocol as the MRI contrast allowed – without subdividing the cortex. Then, we used SmartInterpol [5] to complete the segmentation of the missing slices. Next, we manually corrected the SmartInterpol output as needed, until we were satisfied with the 200 μm isotropic segmentation. The cortex was subdivided using standard FreeSurfer routines. This labelling scheme led to a ground truth segmentation with 98 ROIs, which we have made publicly available (details under “Data Availability”). Supplementary Videos 3 and 4 fly over the coronal and axial slices of the labelled scan, respectively.

As explained in the Results section, we used a simplified version of the *NextBrain* atlas when segmenting the 100 μm scan, in order to better match the ROIs of the automated segmentation and the ground truth (especially in the brainstem). This version was created by replacing the brainstem labels in the histological 3D reconstruction (Fig. 2G, right) by new segmentations made directly in the underlying MRI scan. These segmentations were made with the same methods as for the 100 μm isotropic scan. The new combined segmentations were used to rebuild the atlas.

#### Automated segmentation with Allen MNI template

Automated labelling with the Allen MNI template relied on registration-based segmentation with the NiftyReg package [65,97], which yields state-of-the art performance in brain MRI registration [103]. We used the same deformation model and parameters as the NiftyReg authors used in their own registration-based segmentation work [104]: *(i)* symmetric registration with a deformation model parameterised by a grid of control points (spacing: 2.5 mm = 5 voxels) and B-spline interpolation; *(ii)* local normalised cross correlation as objective function (standard deviation: 2.5mm); and *(iii)* bending energy regularisation (relative weight: 0.001).

#### Linear discriminant analysis (LDA) for AD classification

Linear classification of AD vs controls based on ROI volumes was performed as follows. Leaving one subject out at the time, we used all other subjects to: *(i)* compute linear regression coefficients to correct for sex and age (intracranial volume was corrected by division); *(ii)* estimate mean vectors for the two classes 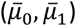, as well as a pooled covariance matrix (Σ); and *(iii)* use the means and covariance to compute an unbiased log-likehood criterion *L* for the left-out subject:

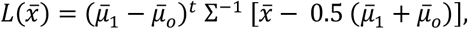

where 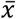 is the vector with ICV-, sex-, and age-corrected volumes for the left-out subject. Once the criterion *L* has been computed for all subjects, we it can be globally thresholded for accuracy and ROC analysis. We note that, for *NextBrain*, the high number of ROIs renders the covariance matrix singular. We prevent this by using regularised LDA: we normalise all the ROIs to unit variance and then compute the covariance as Σ = S + λI, where S is the sample covariance, *I* is the identity matrix, and λ = 1.0 is a constant. We note that normalizing to unit variance enables us to use a fixed, unit λ – rather than having to estimate λ for every left-out subject.

#### B-spline fitting of aging trajectories

To compute the B-spline fits in Extended Data Fig. 8, we first corrected the ROI volumes by sex (using regression) and intracranial volume (by division). Next, we modelled the data with a Laplace distribution, which is robust against outliers which may be caused by potential segmentation mistakes. Specifically, we used an age-dependent Laplacian where the location *μ* and scale *b* are both B-splines with four evenly space control points at 30, 51.6, 73.3, and 95 years. The fit is optimised with gradient ascent over the log-likelihood function:

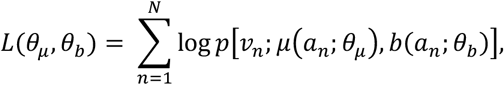

where *p*(*x; μ, b*) is the Laplace distribution with location *μ* and scale *b*; *v*_*n*_ is the volume of ROI for subject *n*; *a*_*n*_ is the age of subject *n*; *μ*(*a*_*n*_; *θ* _*μ*_) is a B-spline describing the location, parameterised by *θ;* and *b*(*a*_*n*_; *θ*_*b*_) is a B-spline describing the scale, parameterised by *θ*. The 95% confidence interval of the Laplace distribution is given by *μ* ± 3*b*.

## Supporting information

Supplementary Atlas Ontology

Supplementary comparison to other atlases

## Acknowledgements

The authors would like to thank the donors, without whom this work would have not been possible. The authors would also like to acknowledge Dr. Paul Johns for his invaluable courses in neuroanatomy at St George’s, University of London.

## Author contributions

- *Conceptualization:* JCA, BLE, JLH, ZJ, JEI
- *Data curation:* AC, MM, ER, JA, SC, EB, BB, AA, LZ, DLT, DK, MB
- *Formal analysis:* AC, LZ, JEI
- *Funding acquisition:* JEI
- *Investigation:* AC, MM, ER, LP, RA, JA, SC, EB, LZ, JEI
- *Methodology:* AC, MM, OP, YB, JLH, ZJ, JEI
- *Project administration:* ER, JLH, CS, ZJ, JEI
- *Resources:* DLT, JLH, CS, ZJ
- *Software:* AC, MM, BB, AA, OP, YB, PS, JH, JEI
- *Supervision:* ER, LP, DK, MB, JLH, CS, ZJ, JEI
- *Validation:* AC, JEI
- *Visualisation:* AC, PS, JH, JEI
- *Writing – original draft:* AC, ER, JEI
- *Writing – review & editing:* all authors.

## Funding

This research has been primarily funded by the European Research Council awarded to JEI (Starting Grant 677697, project “BUNGEE-TOOLS”). MB is supported by a Fellowship award from the Alzheimer’s Society, UK (AS-JF-19a-004-517). OP is supported by a grant from the Lundbeck foundation (R360–2021–39). MM is supported by the Italian National Institute of Health with a Starting Grant, and by the Wellcome Trust through a Sir Henry Wellcome Fellowship (213722/Z/18/Z). Further support was provided by NIH grants 1RF1MH123195, 1R01AG070988, 1UM1MH130981, and 1RF1AG080371.

## Data availability

The raw data used in this article (MRI, histology, segmentations, etc.) can be downloaded from: https://doi.org/10.5522/04/24243835

An online tool to interactively explore the 3D reconstructed data can be found here: https://github-pages.ucl.ac.uk/NextBrain

This website also includes links to videos, publications, code, and other resources.

The segmentation of the *ex vivo* scan can be found at: https://openneuro.org/datasets/ds005422/versions/1.0.1

## Code availability

The code used in this article for 3D histology reconstruction can be downloaded from: https://github.com/acasamitjana/ERC_reconstruction and used and distributed freely.

The segmentation tool is provided as Python code and is integrated in our neuroimaging toolkit “FreeSurfer”: https://surfer.nmr.mgh.harvard.edu/fswiki/HistoAtlasSegmentation. The source code is available on GitHub: https://github.com/freesurfer/freesurfer/tree/dev/mri_histo_util

## Competing interests

The authors have no relevant financial or non-financial interests to disclose.

## Extended Data

**Extended Data Fig. 1:**
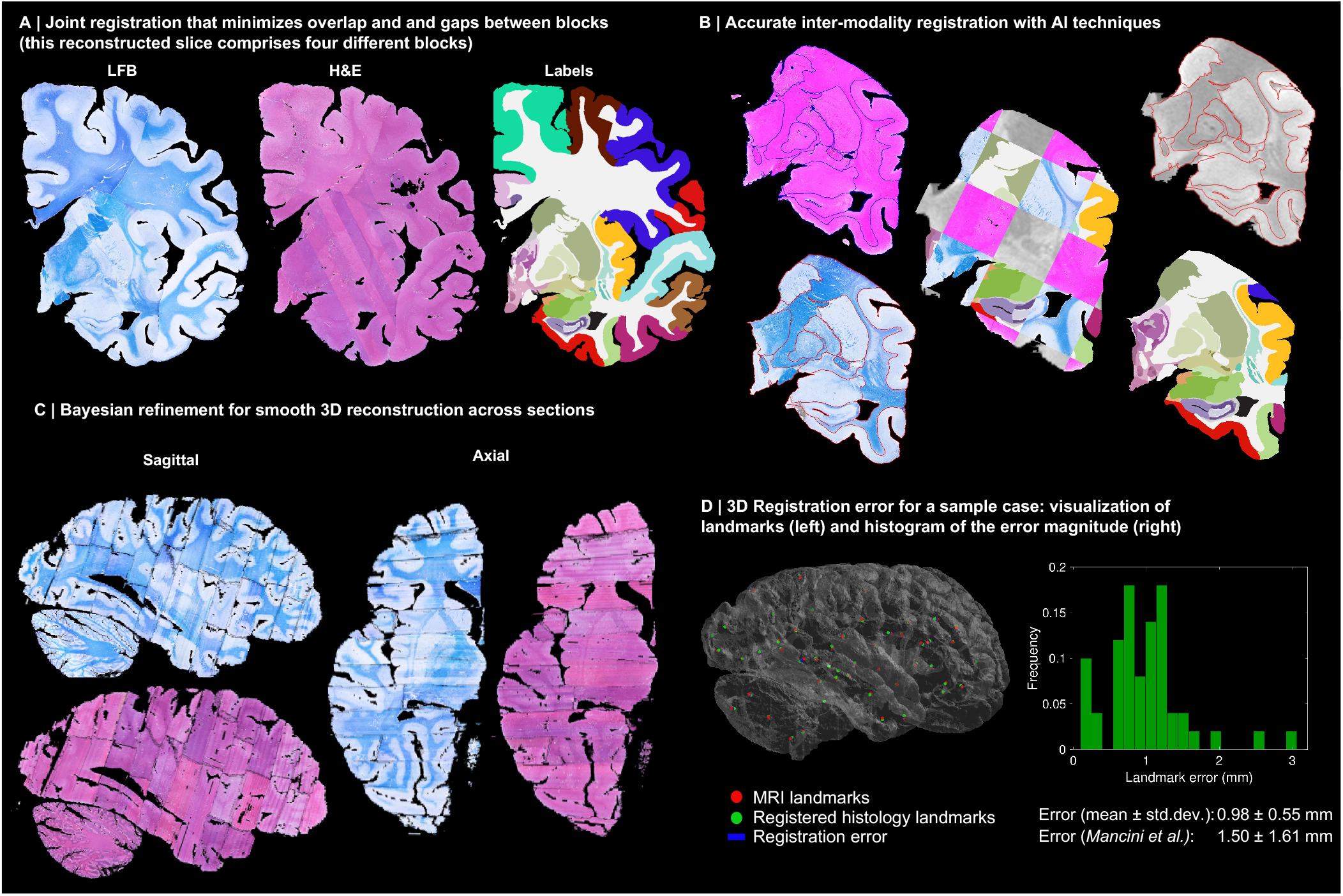
3D reconstruction of Case 2. The visualisation follows the same convention as in Figure 3: (A) Coronal slice of the 3D reconstruction. (B) Registered MRI, LFB, and H&E histology of a block, with tissue boundaries (traced on LFB) overlaid. (C) Orthogonal view of reconstruction, which is smooth and avoids gaps and overlaps. (D) Visualization of 3D landmark registration error (left); histogram of its error (right); and mean ± standard deviation (bottom), compared with our previous pipeline (Mancini et al. [6]).

**Extended Data Fig. 2:**
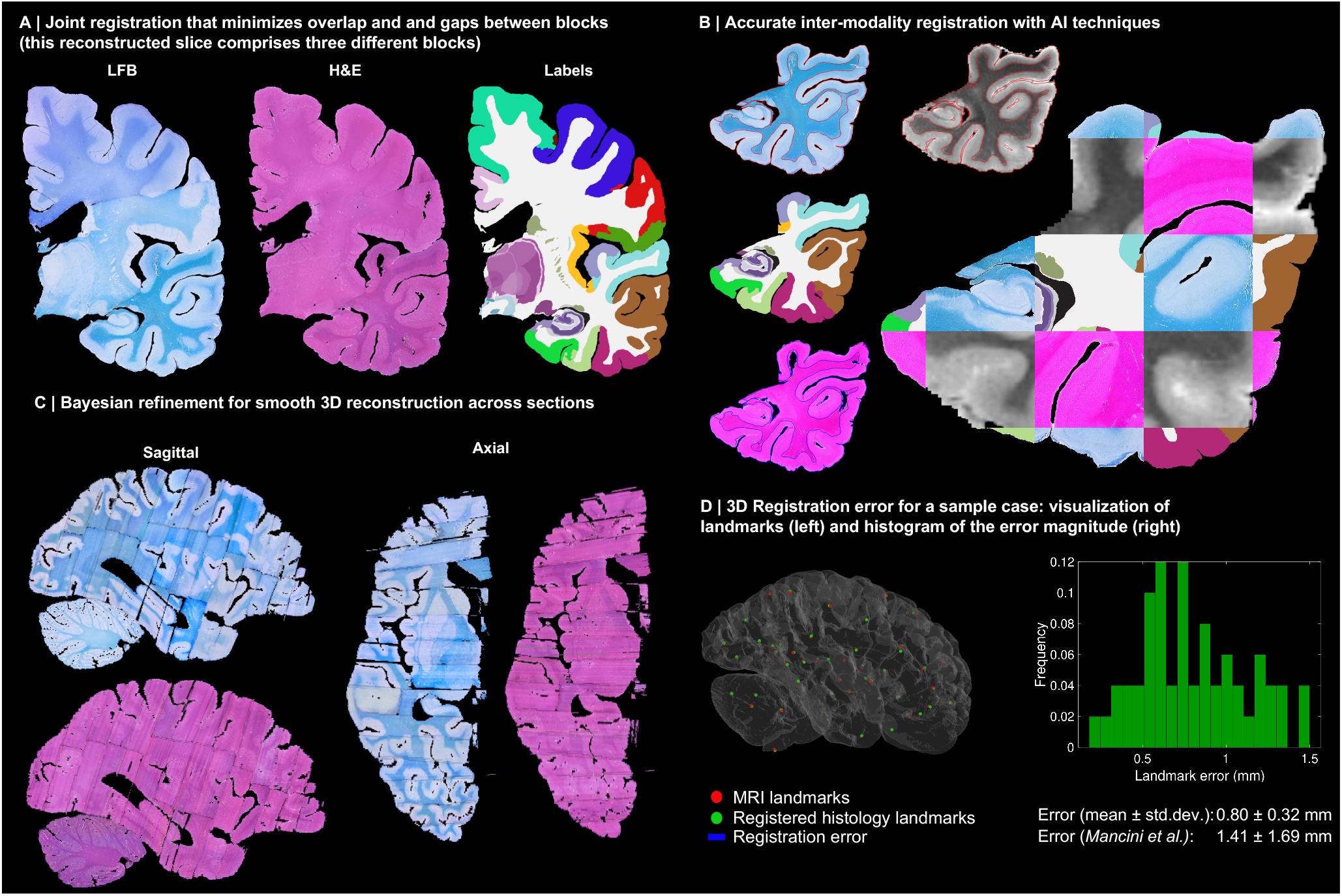
3D reconstruction of Case 3. The visualisation follows the same convention as in Figure 3: (A) Coronal slice of the 3D reconstruction. (B) Registered MRI, LFB, and H&E histology of a block, with tissue boundaries (traced on LFB) overlaid. (C) Orthogonal view of reconstruction, which is smooth and avoids gaps and overlaps. (D) Visualization of 3D landmark registration error (left); histogram of its error (right); and mean ± standard deviation (bottom), compared with our previous pipeline (Mancini et al. [6]).

**Extended Data Fig. 3:**
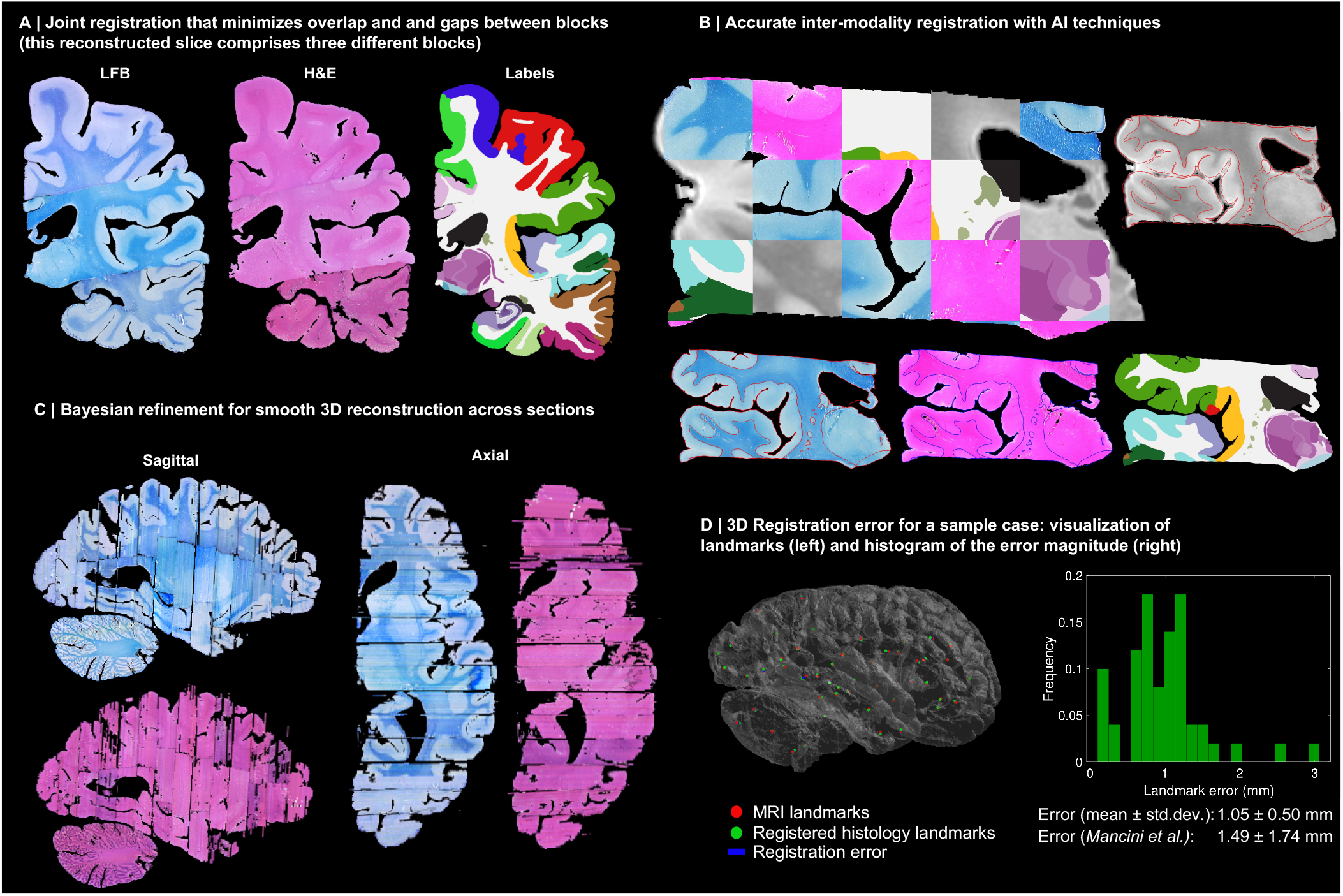
3D reconstruction of Case 4. The visualisation follows the same convention as in Figure 3: (A) Coronal slice of the 3D reconstruction. (B) Registered MRI, LFB, and H&E histology of a block, with tissue boundaries (traced on LFB) overlaid. (C) Orthogonal view of reconstruction, which is smooth and avoids gaps and overlaps. (D) Visualization of 3D landmark registration error (left); histogram of its error (right); and mean ± standard deviation (bottom), compared with our previous pipeline (Mancini et al. [6]).

**Extended Data Fig. 4:**
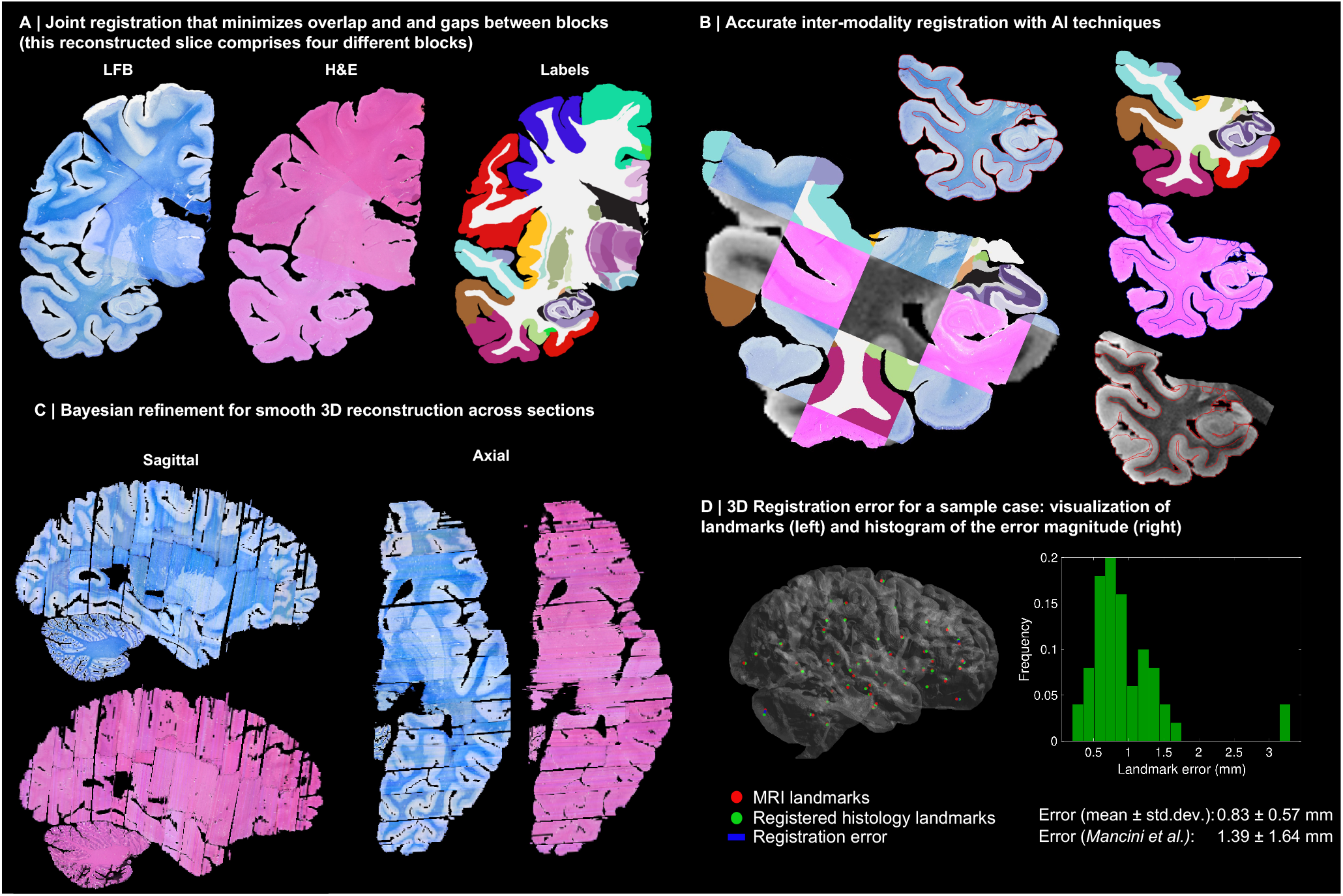
3D reconstruction of Case 5. The visualisation follows the same convention as in Figure 3: (A) Coronal slice of the 3D reconstruction. (B) Registered MRI, LFB, and H&E histology of a block, with tissue boundaries (traced on LFB) overlaid. (C) Orthogonal view of reconstruction, which is smooth and avoids gaps and overlaps. (D) Visualization of 3D landmark registration error (left); histogram of its error (right); and mean ± standard deviation (bottom), compared with our previous pipeline (Mancini et al. [6]).

**Extended Data Fig. 5:**
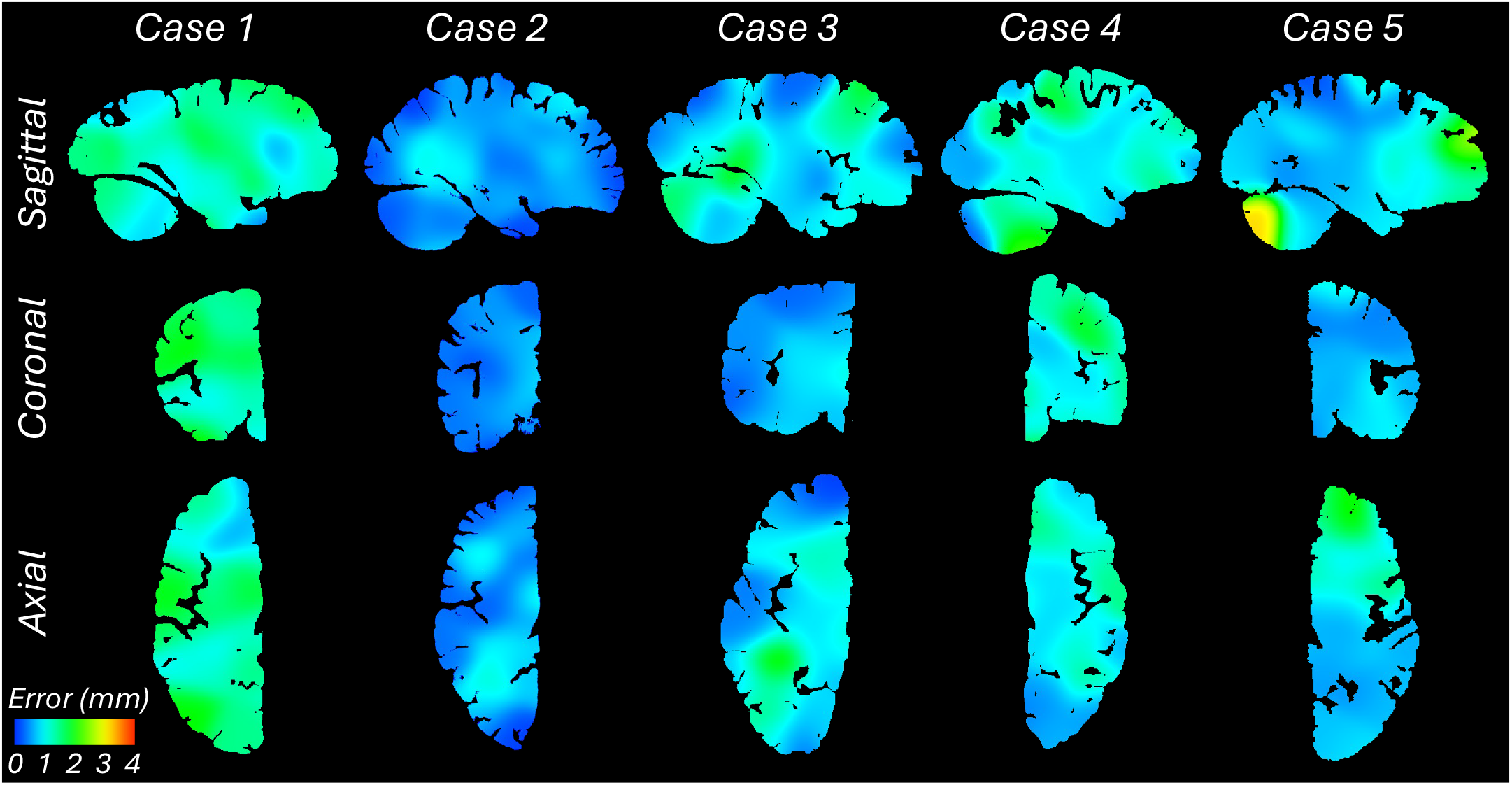
Sagittal, coronal, and axial slices of the continuous maps of the 3D landmark registration error. The maps are computed from the discrete landmarks (displayed in Fig. 3D and Extended Data Figs. 1-4D) using Gaussian kernel regression with σ = 10 mm. There is no clear spatial pattern for the anatomical distribution of the error across subjects.

**Extended Data Tab. 1:**
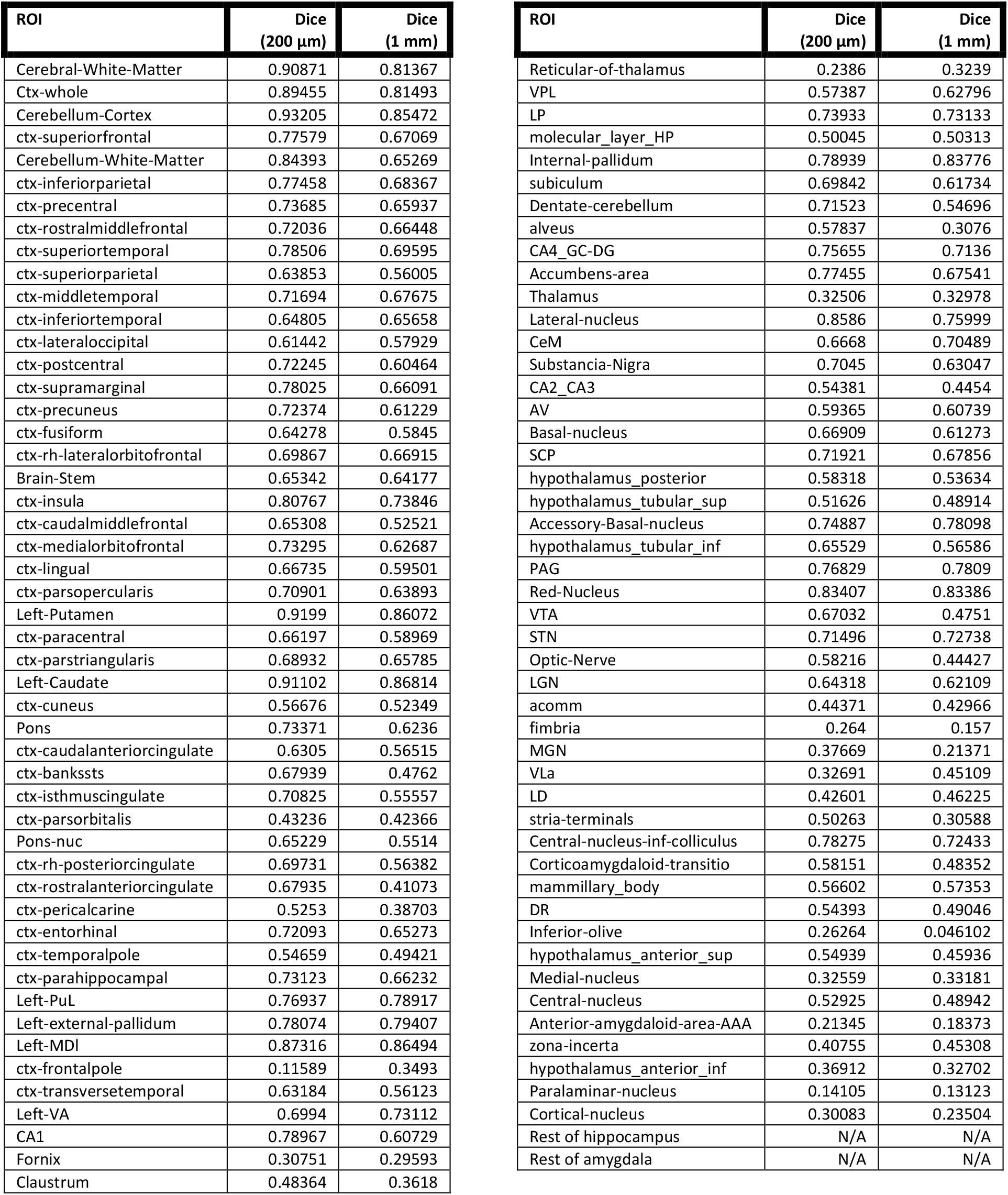
Dice scores between the ground truth labels of the 100 μm *ex vivo* brain MRI scan presented in [3] and the automated segmentations obtained with *NextBrain*. ROIs are listed in decreasing order of size (volume). The Dice scores are shown for segmentations obtained at two different resolutions: 200 μm (the resolution at which we created the ground truth labels) and 1 mm (which is representative of *in vivo* data). We note that the Dice scores are computed from labels made on the right hemisphere (since we did not label the left side of the brain). We also note that the labels “rest of hippocampus” and “rest of amygdala” correspond to voxels that did not clearly belong to any of the manually labelled nuclei, and have therefore no direct correspondence with ROIs in *NextBrain*.

**Extended Data Fig. 6:**
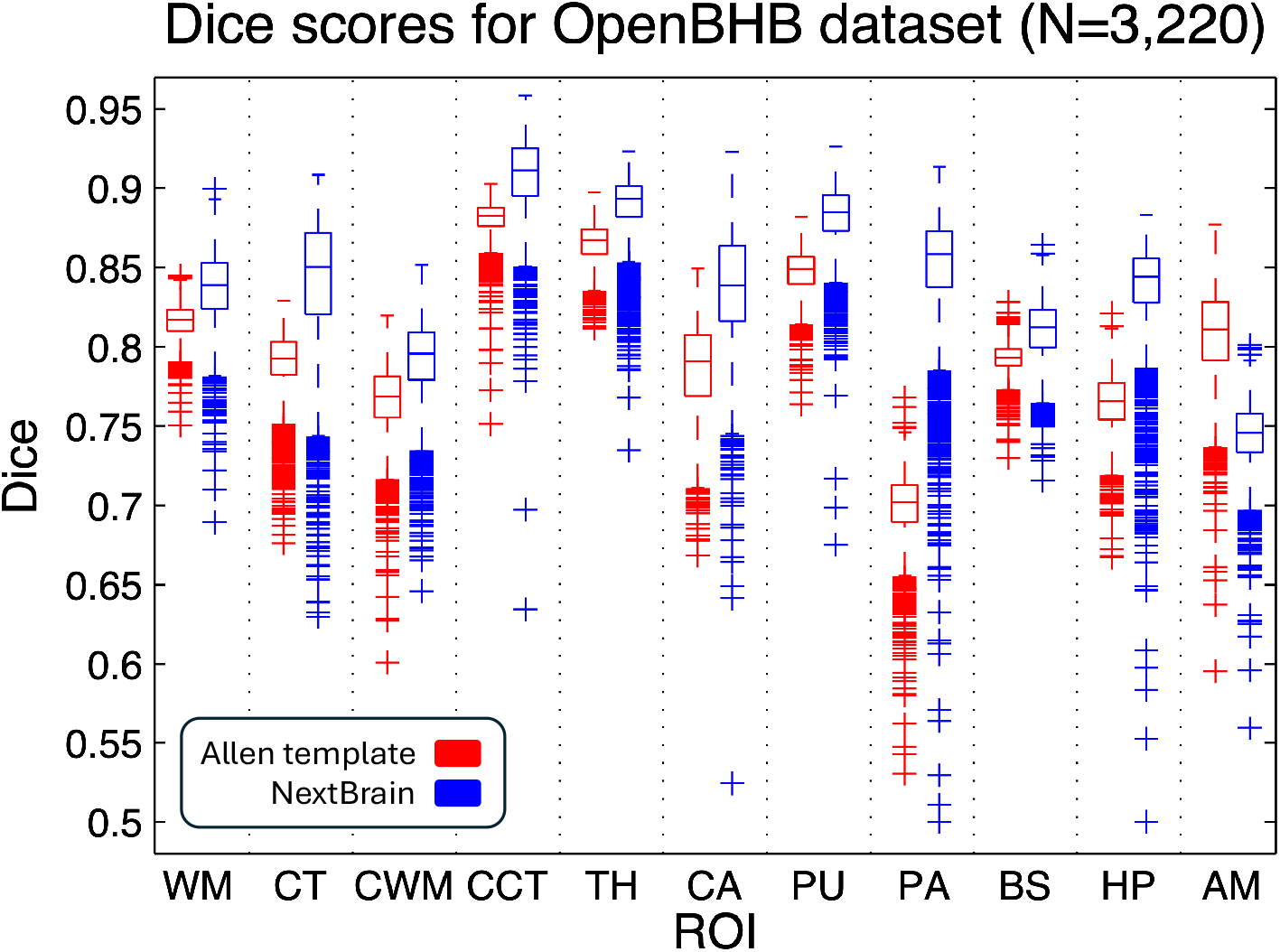
Box plots of the Dice scores for 11 representative ROIs computed on the OpenBHB dataset (3,330 subjects), using the Allen MNI template and NextBrain, with FreeSurfer segmentations as reference. The scores are computed at the whole regions level, i.e., the level of granularity at which FreeSurfer segments. On each box, the central mark indicates the median, the edges of the box indicate the 25_th_ and 75th percentiles, the whiskers extend to the most extreme data points not considered outliers, and the outliers are plotted individually as ‘+’. The abbreviations for the regions are: WM = white matter of the cerebrum, CT = cortex of the cerebrum, CWM = cerebellar white matter, CCT = cerebellar cortex, TH = thalamus, CA = caudate, PU = putamen, PA = pallidum, BS = brainstem, HP = hippocampus, AM = amygdala.

**Extended Data Fig. 7:**
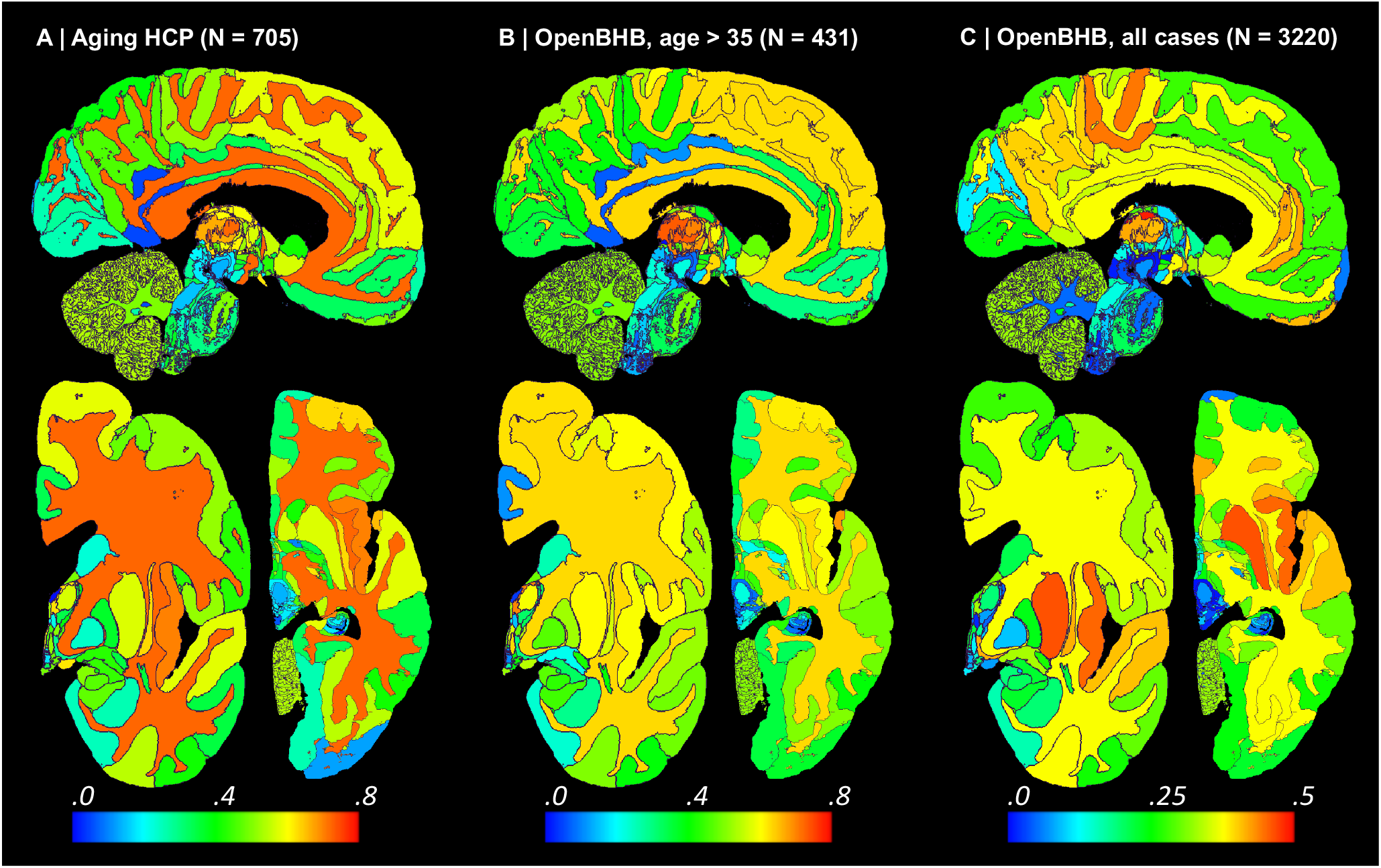
Absolute value of Spearman correlation for ROI volumes vs age derived from in vivo MRI scans (additional slices). The visualisation follows the same convention as in Figure 5: (A) Ageing HCP dataset. (B) OpenBHB dataset, restricted to ages over 35. (C) Full OpenBHB dataset.

**Extended Data Fig. 8:**
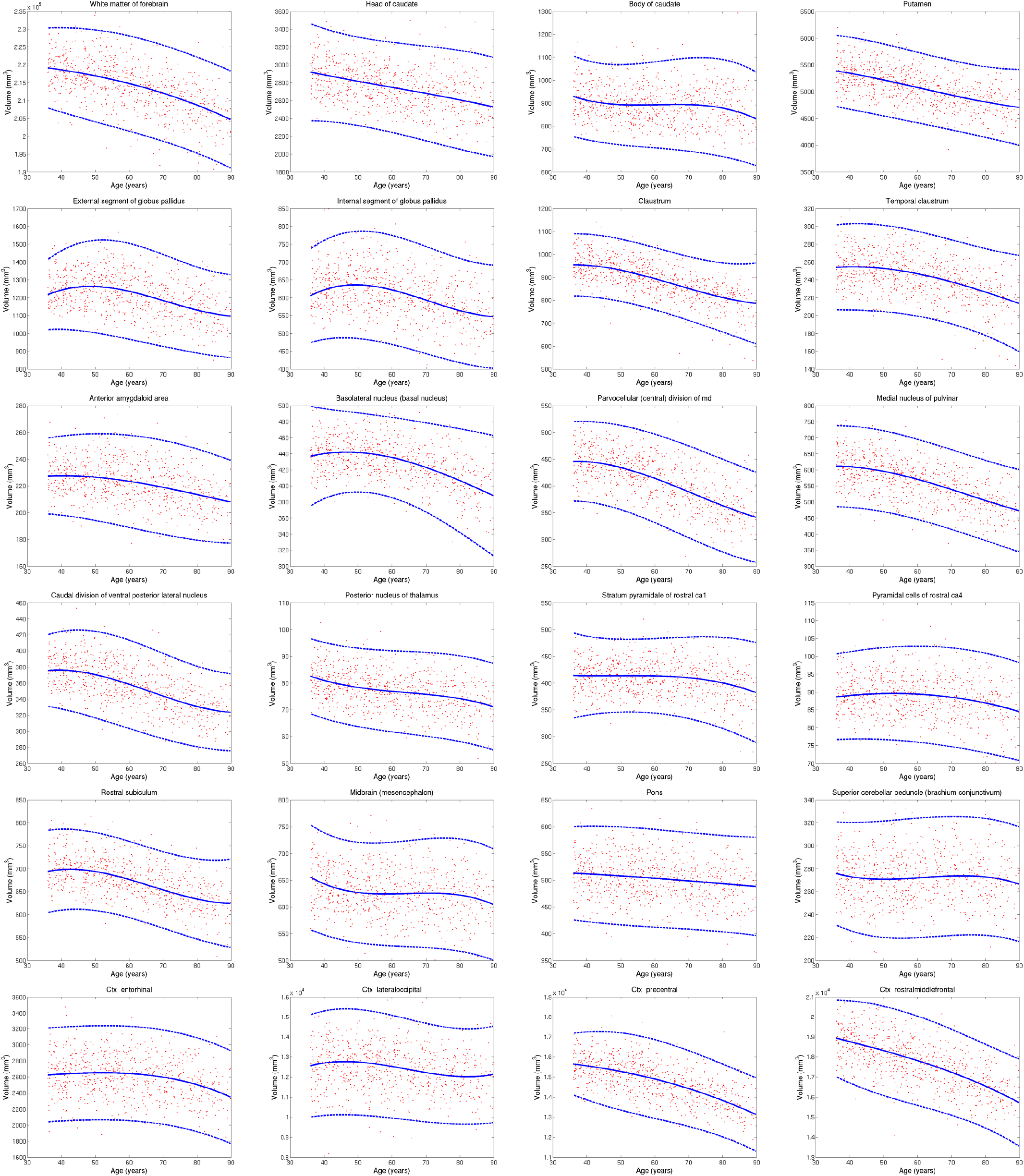
Aging trajectories for select ROIs in HCP dataset, showing differential pattens in subregions of brain structures (thalamus, hippocampus, cortex, etc). The red dots correspond to the ROI volumes of individual subjects, corrected by intracranial volume (by division) and sex (by regression). The blue lines represent the maximum likehood fit of a Laplace distribution with location and scale parameters parametrised by a B-spline with four control points (equally space between 30 and 95 years). The continuous blue line represents the location, whereas the dashed lines represent the 95% confidence interval (equal to three times the scale parameter in either direction). Volumes of contralateral structures are averaged across left and right.

## Notes

https://github-pages.ucl.ac.uk/NextBrain

### Competing Interest Statement

The authors have declared no competing interest.

### Summary of Updates

New experiments and results, as well as new figures.

