## Supplementary comparison to other atlases for "A next-generation, histological atlas of the human brain and its application to automated brain MRI segmentation"

This supplement contains a qualitative comparison of the Mai-Paxinos atlas, the Allen atlas and our atlas *NextBrain*. Every page in our comparison corresponds to each of the sections that are available in the digital version of the Mai-Paxinos atlas, available at:
https://www.thehumanbrain.info/brain/sections.php

On the left of each page, we show the labels and the histological sections by Mai-Paxinos. On the right, we show the approximately equivalent section (and corresponding labels) of the Allen reference brain, available at:
http://atlas.brain-map.org

In the middle of each page, we show an approximately corresponding coronal slice of *NextBrain* (with the boundaries between the most likely regions overlaid in red). We note that there are anatomical differences due to:

- The slightly different orientation of their sections, compared with *NextBrain*.
- The probabilistic nature of *NextBrain* - which is particularly noticeable in cortical regions.
- Anatomical variability between the single specimens that the atlases are based on – while ours is an average.

In general, the agreement between NextBrain and the two atlases is good, especially for the outer boundaries of the main structures, and for the borders between the grey matter and white matter and/or the cerebrospinal fluid. This is particularly clear in the striatum, the hippocampus, the amygdala and the thalamus. The main differences lie in:

- The delineation of sub-structures within the main cortical regions. For example, the superior frontal gyrus is labelled as a whole in both Allen and *NextBrain*, rather than separating its medial and lateral part as done in the Mai-Paxinos. Similarly, the insula is labelled as a whole in both the Mai-Paxinos atlas and *NextBrain*, rather than into its agranular and dysgranular subcomponents as in the Allen atlas. We note that *NextBrain* directly takes the cortical parcellation from FreeSurfer.
- The annotation of the smaller nuclei within the subcortical regions (e.g., the nucleus accumbens as a whole in NextBrain, rather than its lateral, medial and central part in the Mai-Paxinos or its shell and core in the Allen). We emphasise that these differences are present *even between the Mai-Paxinos and Allen*, and they are mainly due to the forced choice of applying arbitrary anatomical criteria for poor visualization and insufficient contrast in smaller regions, and to different anatomical definitions used by anatomists.
- Increased thickness of some regions (particularly cortical) in *NextBrain* when taking the maximum-probability labels, due to the probabilistic nature of the atlas.

Some further comments about specific brain regions:

- The smaller subnuclei of the hypothalamus are difficult to delineate and accurately replicate (see, e.g., differences between Mai-Paxinos and Allen). *NextBrain* is fairly close to Allen in this area. We also note that larger subregions (anterior-inferior, anterior-superior, tubular-inferior, tubular-superior, and posterior with the mammillary bodies) are can be reliable segmented from in vivo MRI ("Automated segmentation of the hypothalamus and associated subunits in brain MRI." *Neuroimage* 223, 2020: 117287).

- Although the outer boundaries of the hippocampus match between the atlases, there are some differences in the anatomical delineation when it comes to the subfields and internal boundaries. For example, in our atlas the CA3 region includes part of the CA2, and the CA4 the molecular layer of the dentate gyrus, rather than separating them into individual components.

- The overall boundaries and division of the amygdala into its broad subnuclei is generally consistent across the three atlases. One of the main differences between *NextBrain* and the other atlases is the more detailed subdivision of the main nuclei in Mai-Paxinos and Allen. While an extended segmentation of the larger nuclei (e.g., lateral, basal, accessory basal) would be certainly feasible in the future, subdividing smaller nuclei (e.g., medial into rostral and caudal subdivision; cortical into anterior and posterior) will always be challenging to robustly delineate. Again, we emphasise that there are clear differences between the Mai-Paxinos and Allen atlases in these subdivisions, particularly for smaller regions of interest.

- The thalamus has some clear differences between atlases, due to the complexity of visually identifying reliable boundaries between its subnuclei. However, the main subdivisions (e.g., pulvinar or mediodorsal regions) are consistent across the three atlases.

*Max & Paxinos (histology)*

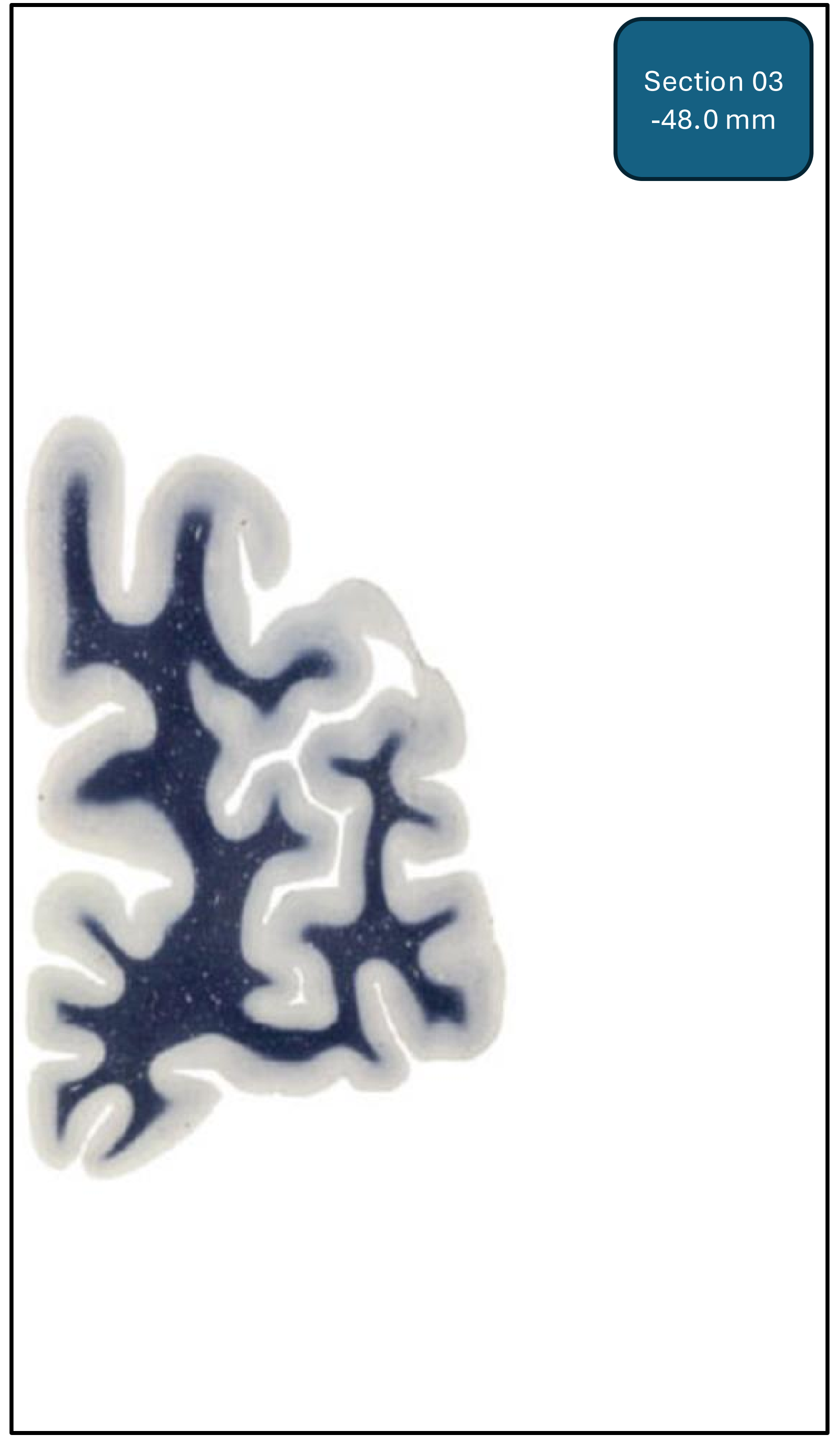


*Max & Paxinos (labels)*

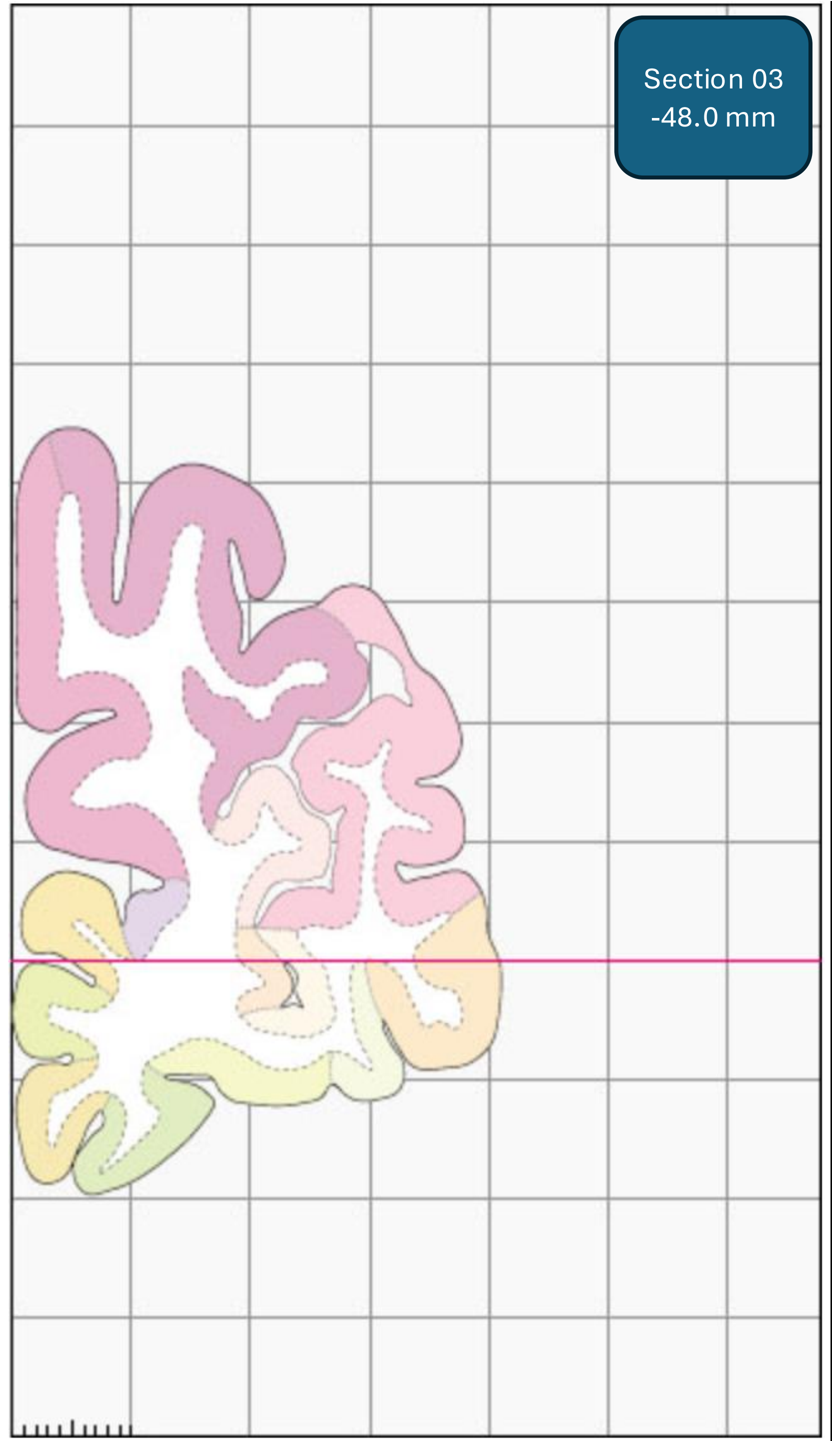


*NextBrain (probabilistic labels)*

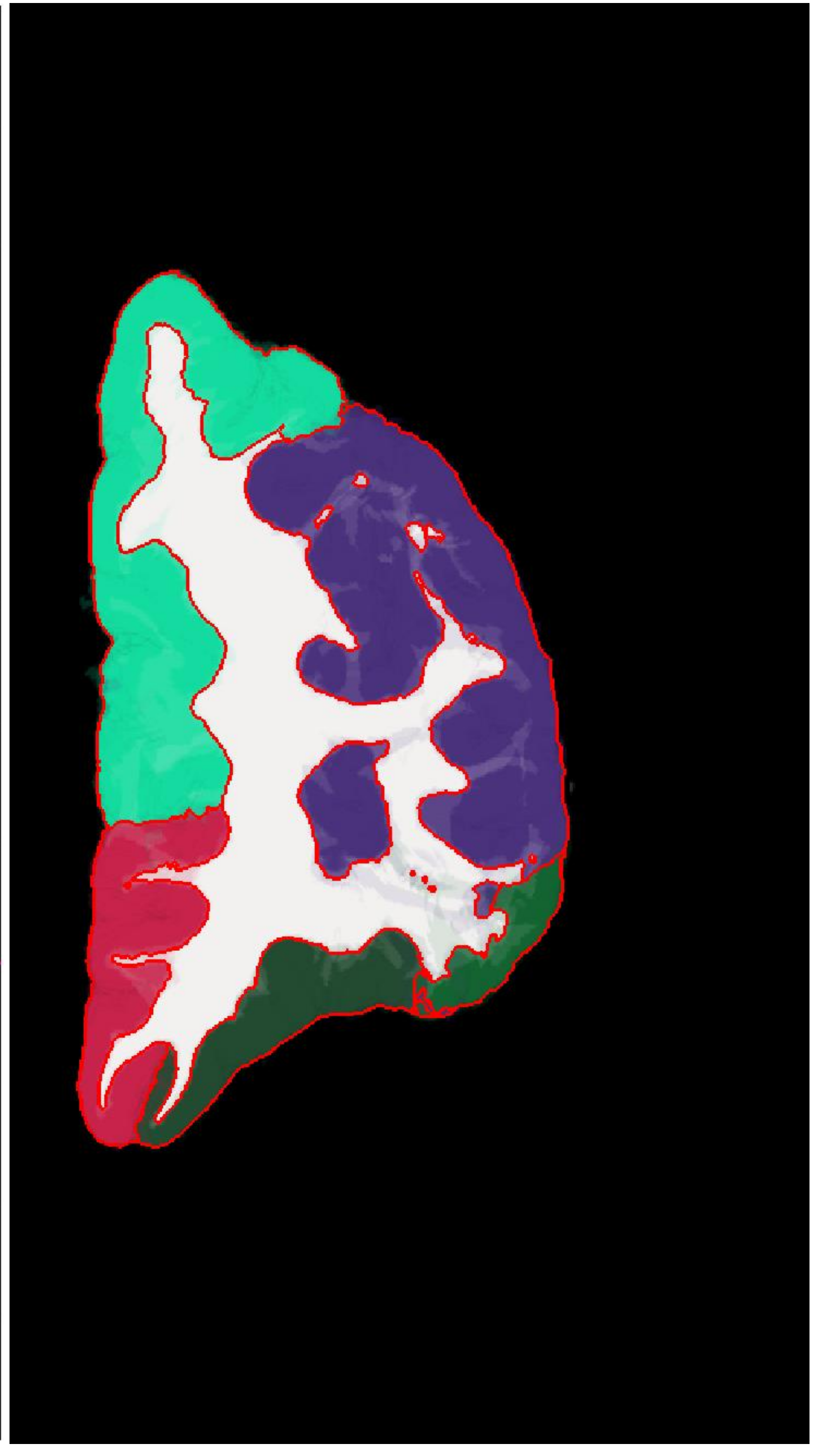

*Allen reference brain (histology)*

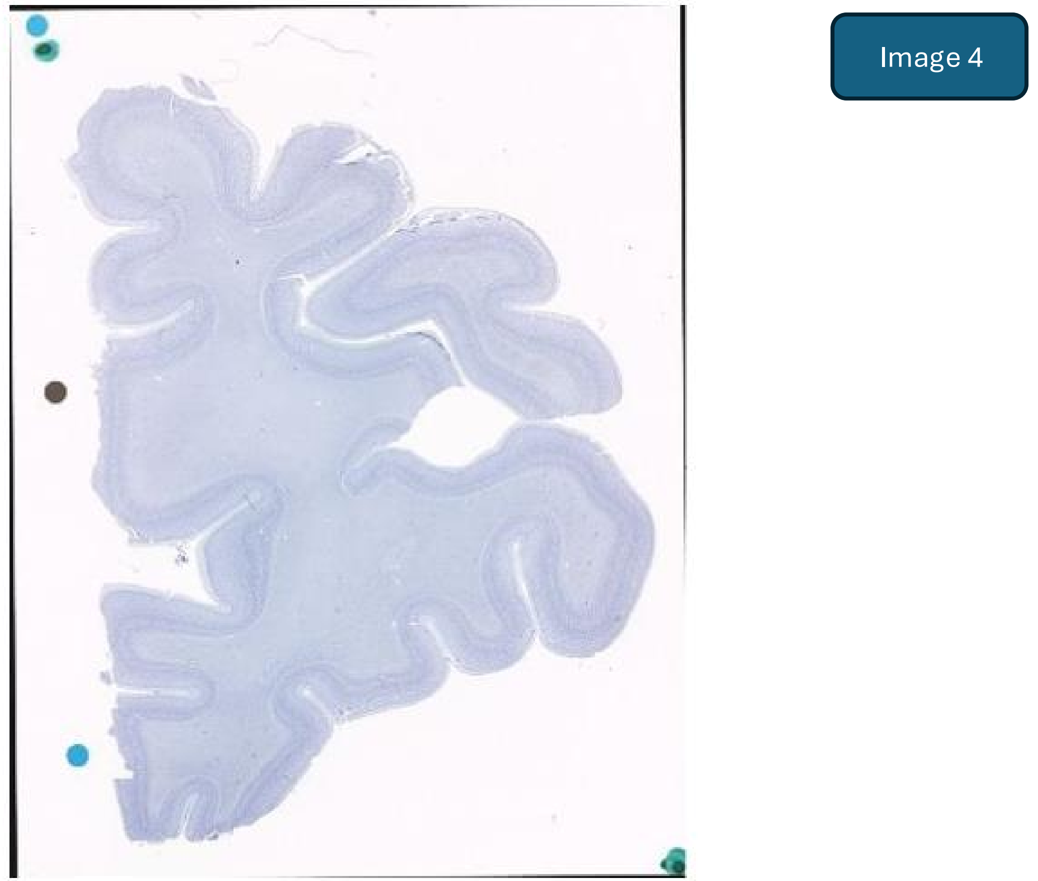


*Allen reference brain (labels)*

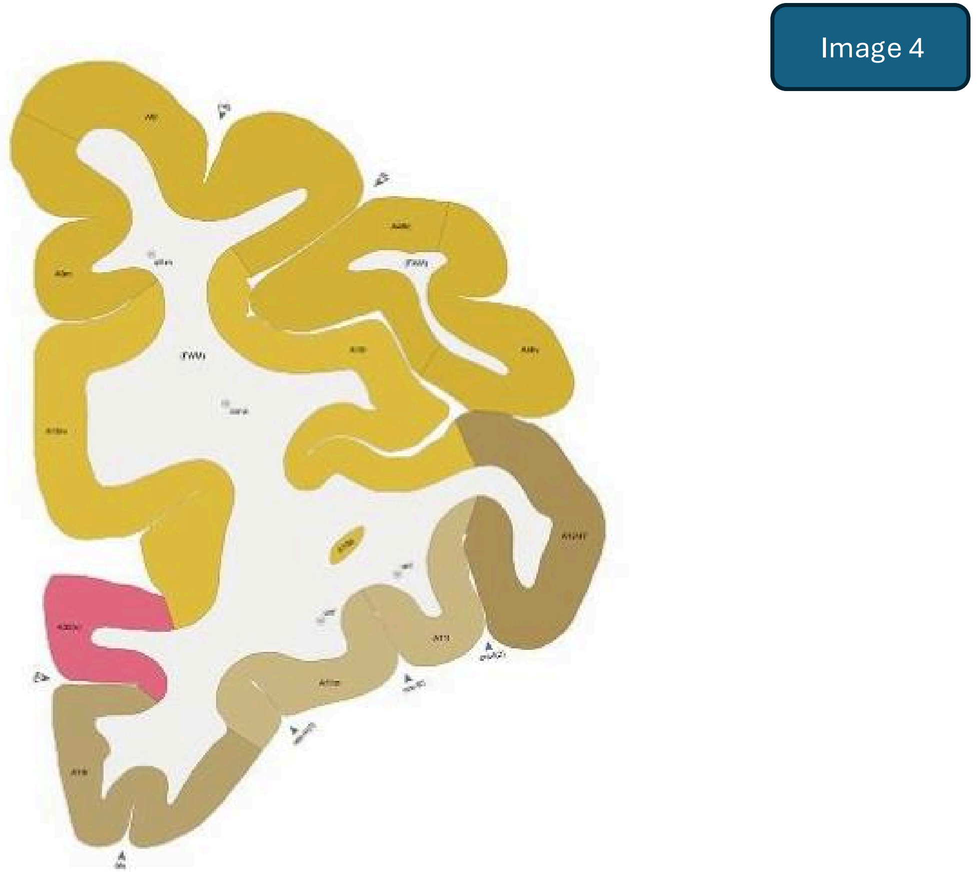

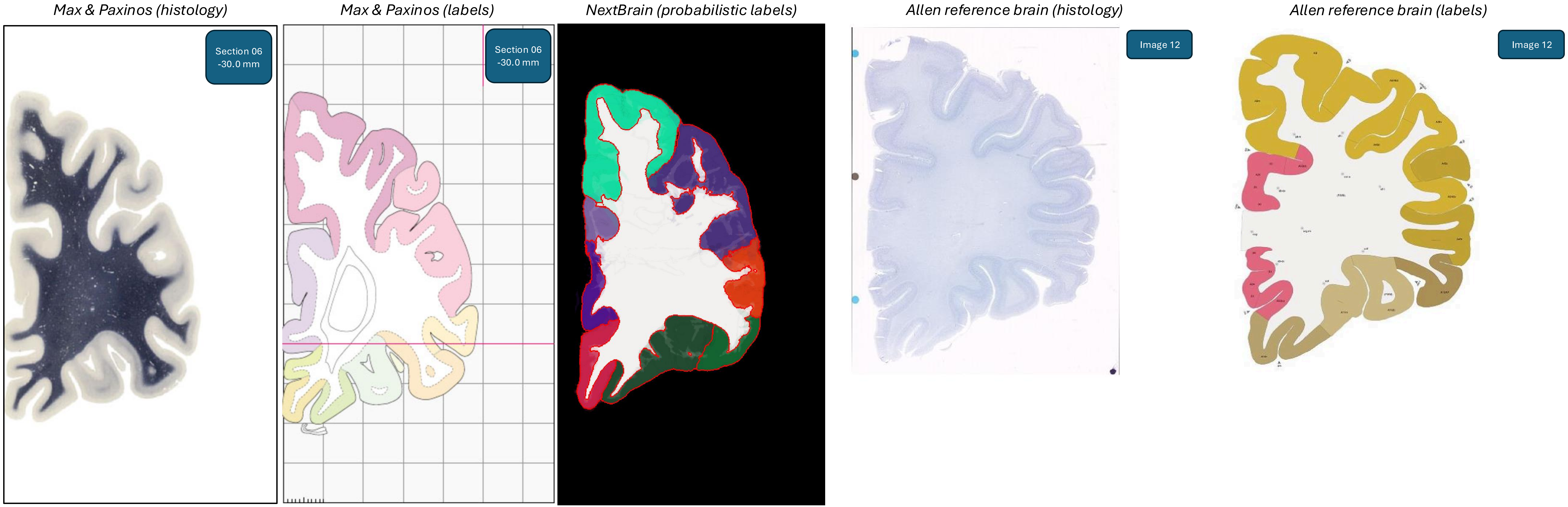
Max & Paxinos (histology)
Section 06
-30.0 mm
Max & Paxinos (labels)
Section 06
-30.0 mm
NextBrain (probabilistic labels)
Allen reference brain (histology)
Image 12
Allen reference brain (labels)
Image 12

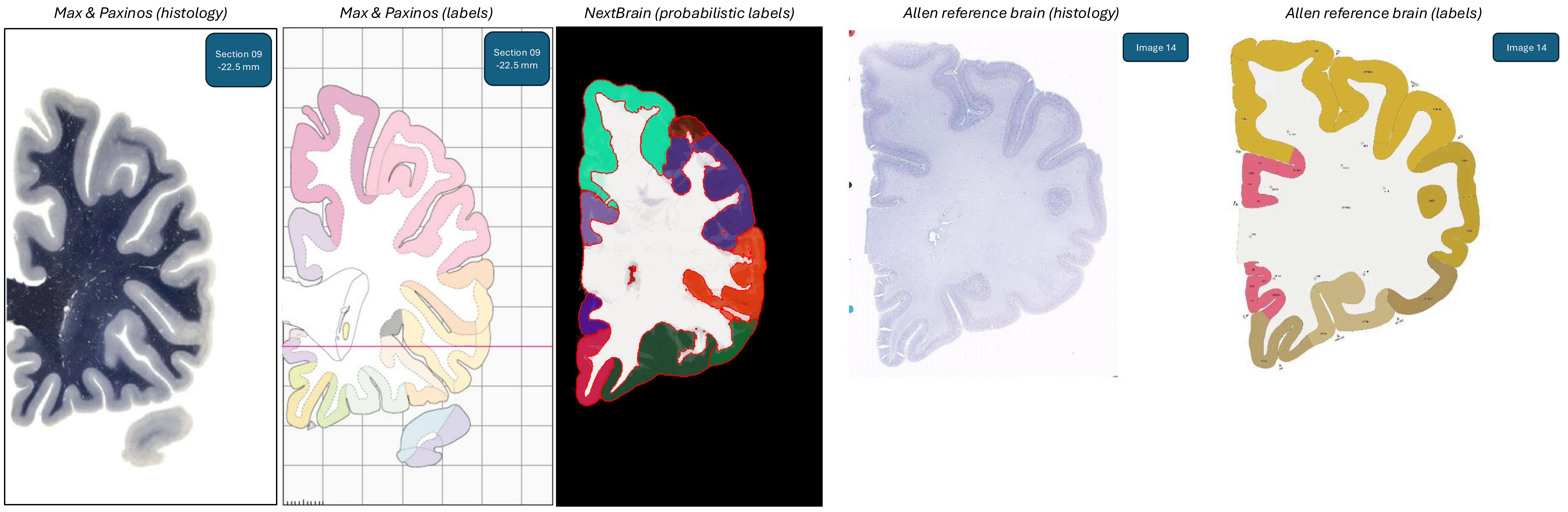
Max & Paxinos (histology)
Section 09
-22.5 mm
Max & Paxinos (labels)
Section 09
-22.5 mm
NextBrain (probabilistic labels)
Allen reference brain (histology)
Image 14
Allen reference brain (labels)
Image 14

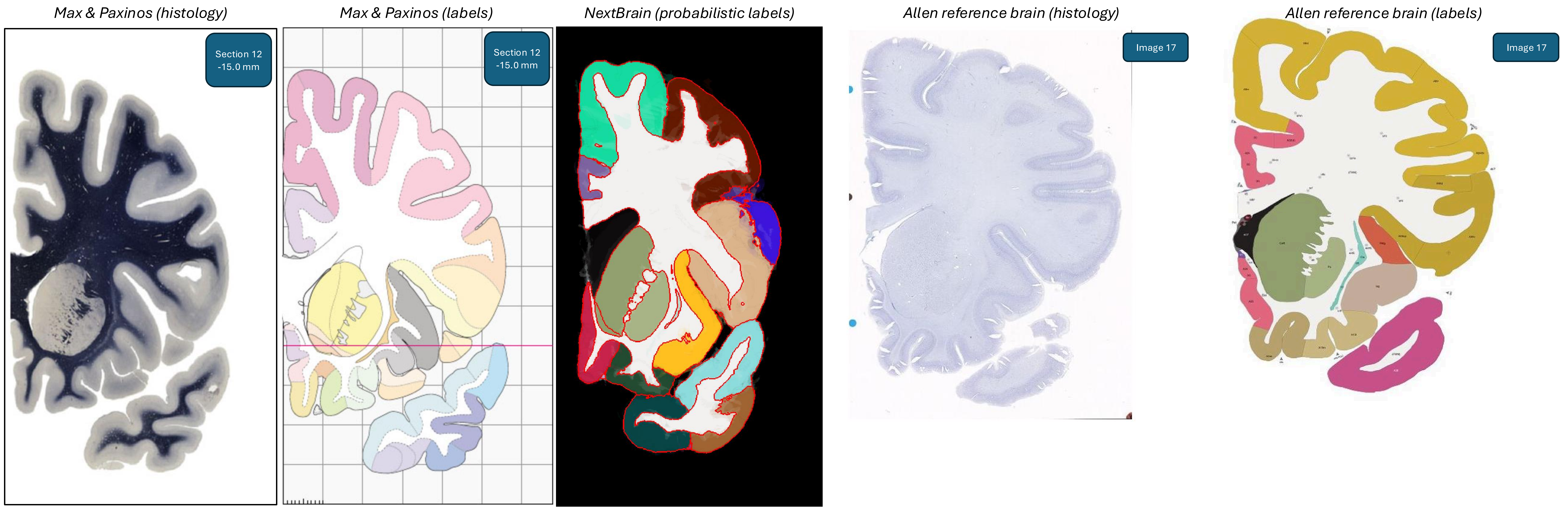
Max & Paxinos (histology)
Section 12
-15.0 mm
Max & Paxinos (labels)
Section 12
-15.0 mm
NextBrain (probabilistic labels)
Allen reference brain (histology)
Image 17
Allen reference brain (labels)
Image 17

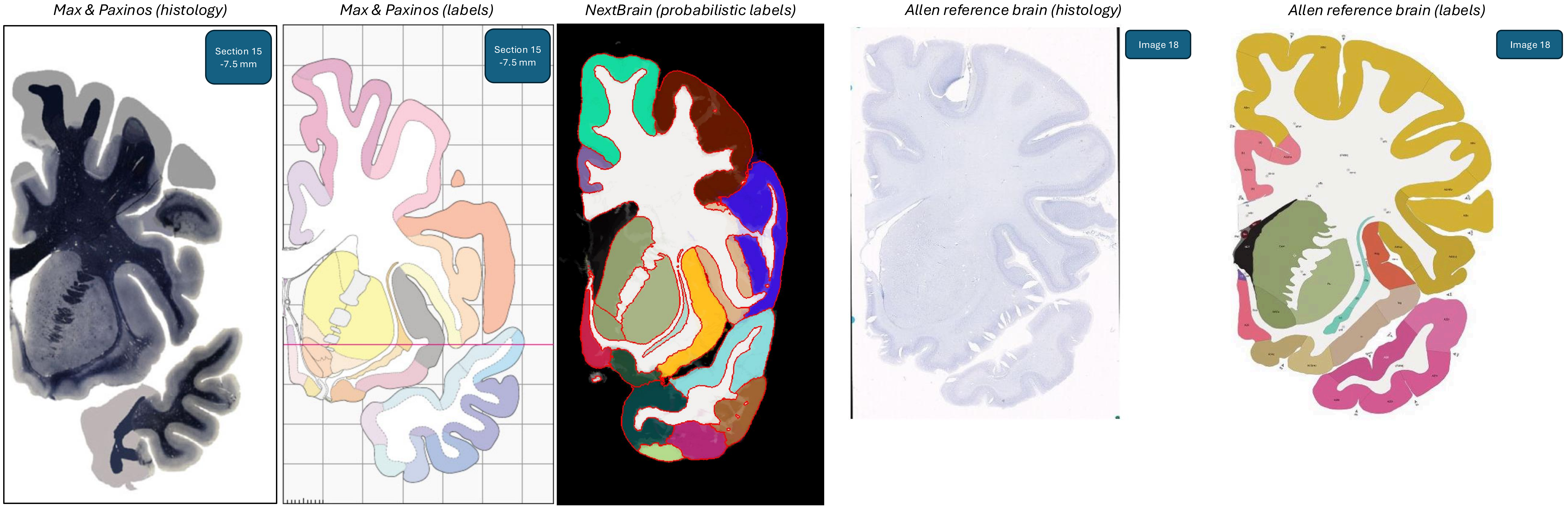
Max & Paxinos (histology)
Section 15
-7.5 mm
Max & Paxinos (labels)
Section 15
-7.5 mm
NextBrain (probabilistic labels)
Allen reference brain (histology)
Image 18
Allen reference brain (labels)
Image 18

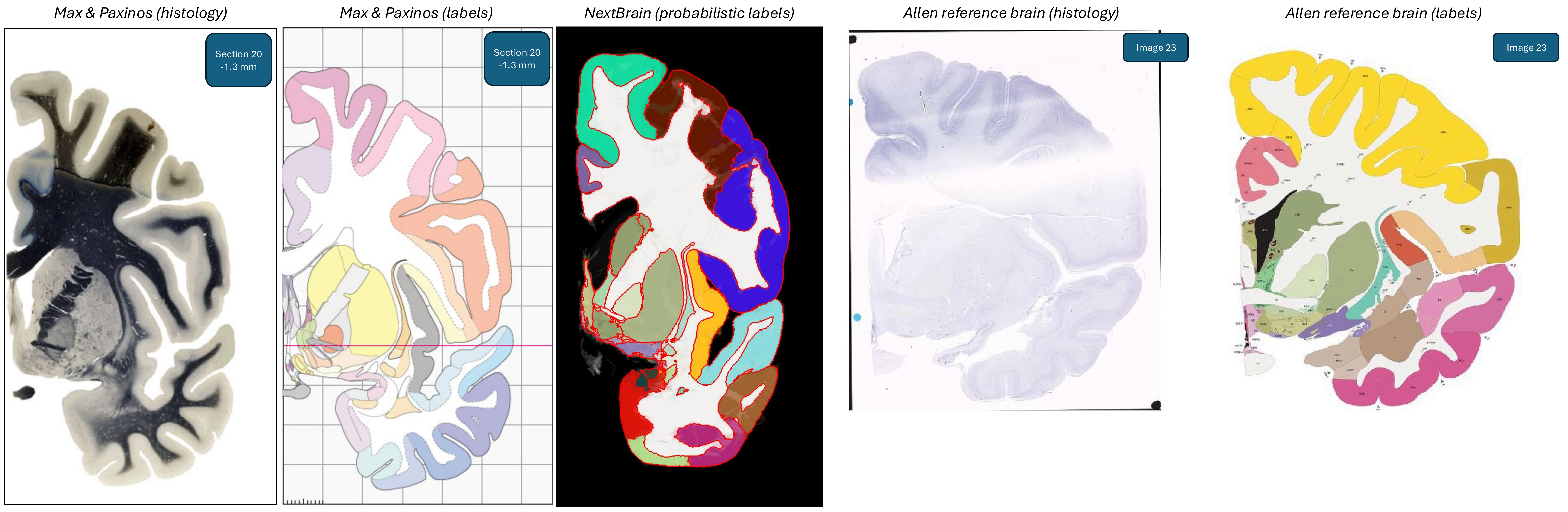
Max & Paxinos (histology)
Section 20
-1.3 mm
Max & Paxinos (labels)
Section 20
-1.3 mm
NextBrain (probabilistic labels)
Allen reference brain (histology)
Image 23
Allen reference brain (labels)
Image 23

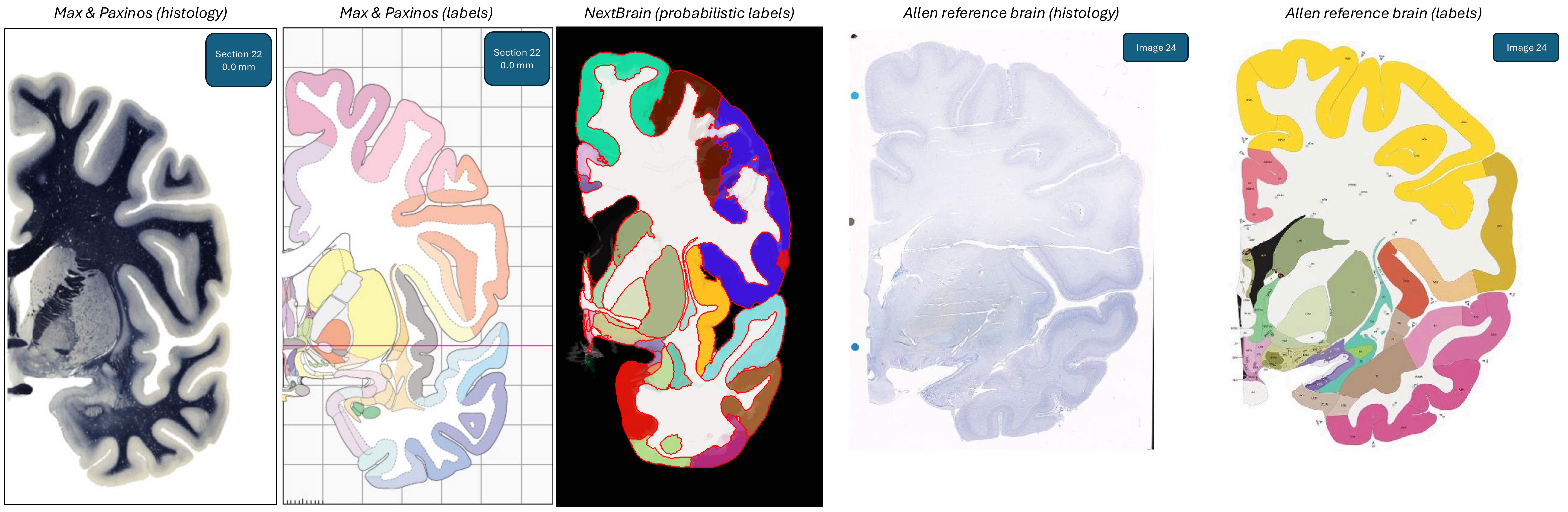
Max & Paxinos (histology)
Max & Paxinos (labels)
NextBrain (probabilistic labels)
Allen reference brain (histology)
Allen reference brain (labels)
Section 22
0.0 mm
Section 22
0.0 mm
Image 24
Image 24

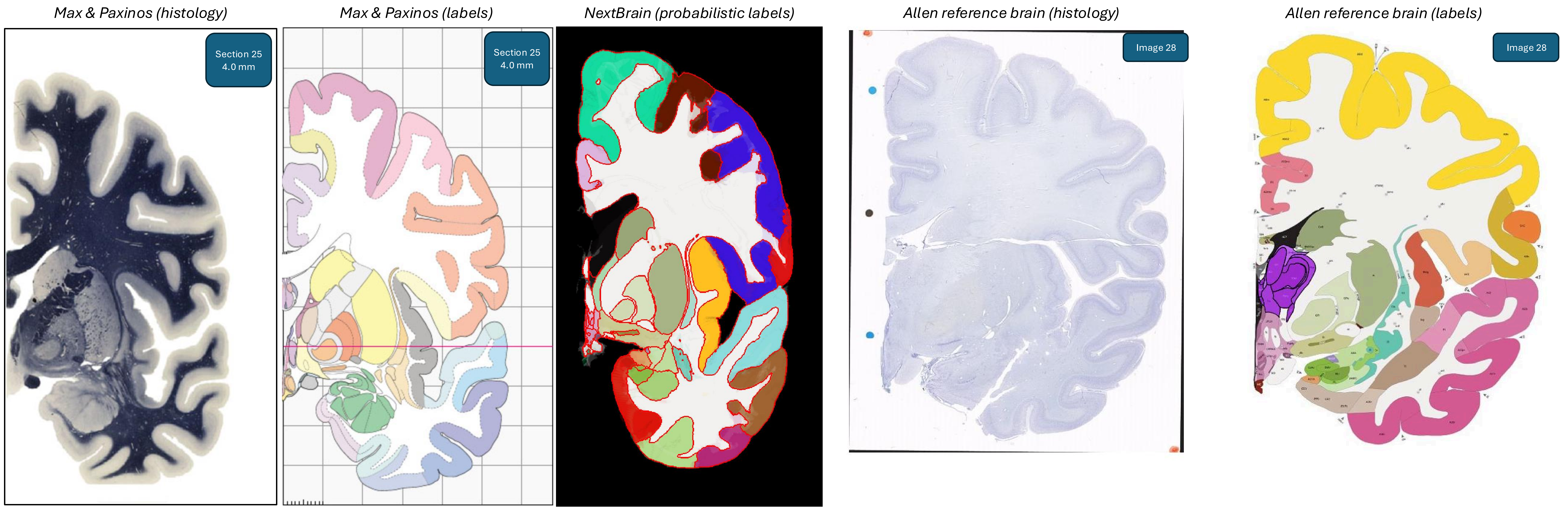

Max & Paxinos (histology)
Section 25
4.0 mm
Max & Paxinos (labels)
Section 25
4.0 mm
NextBrain (probabilistic labels)
Allen reference brain (histology)
Image 28
Allen reference brain (labels)
Image 28

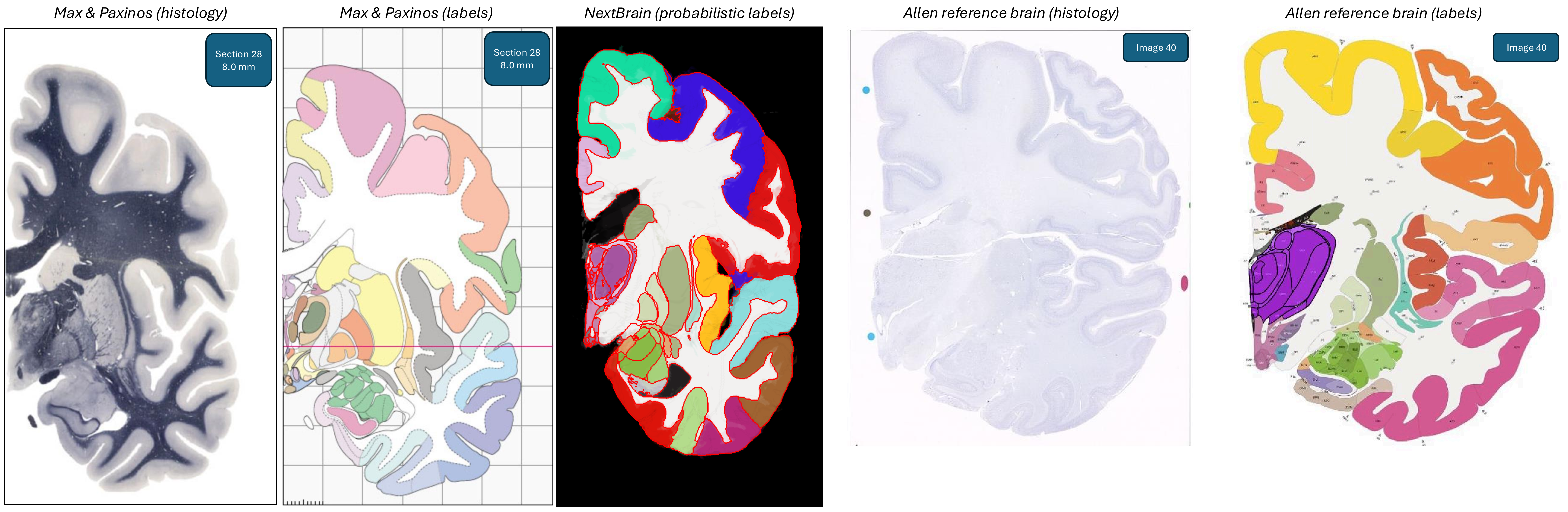
Max & Paxinos (histology)
Section 28
8.0 mm
Max & Paxinos (labels)
Section 28
8.0 mm
NextBrain (probabilistic labels)
Allen reference brain (histology)
Image 40
Allen reference brain (labels)
Image 40

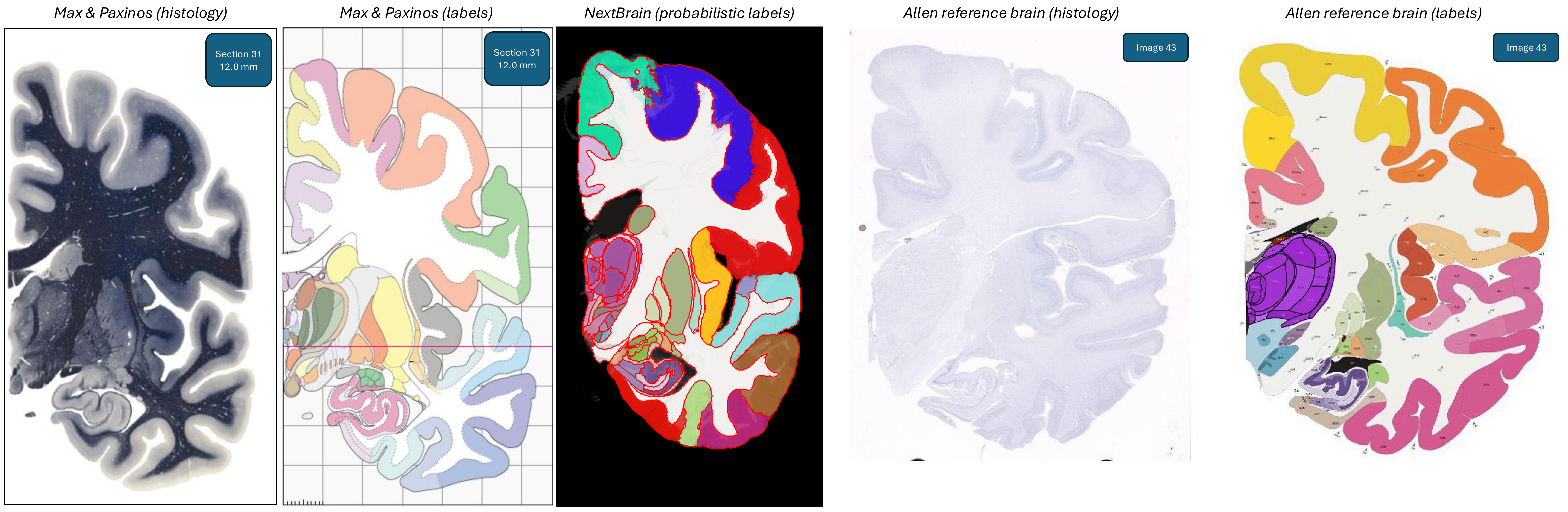
Max & Paxinos (histology)
Max & Paxinos (labels)
NextBrain (probabilistic labels)
Allen reference brain (histology)
Allen reference brain (labels)
Section 31
12.0 mm
Section 31
12.0 mm
Image 43
Image 43

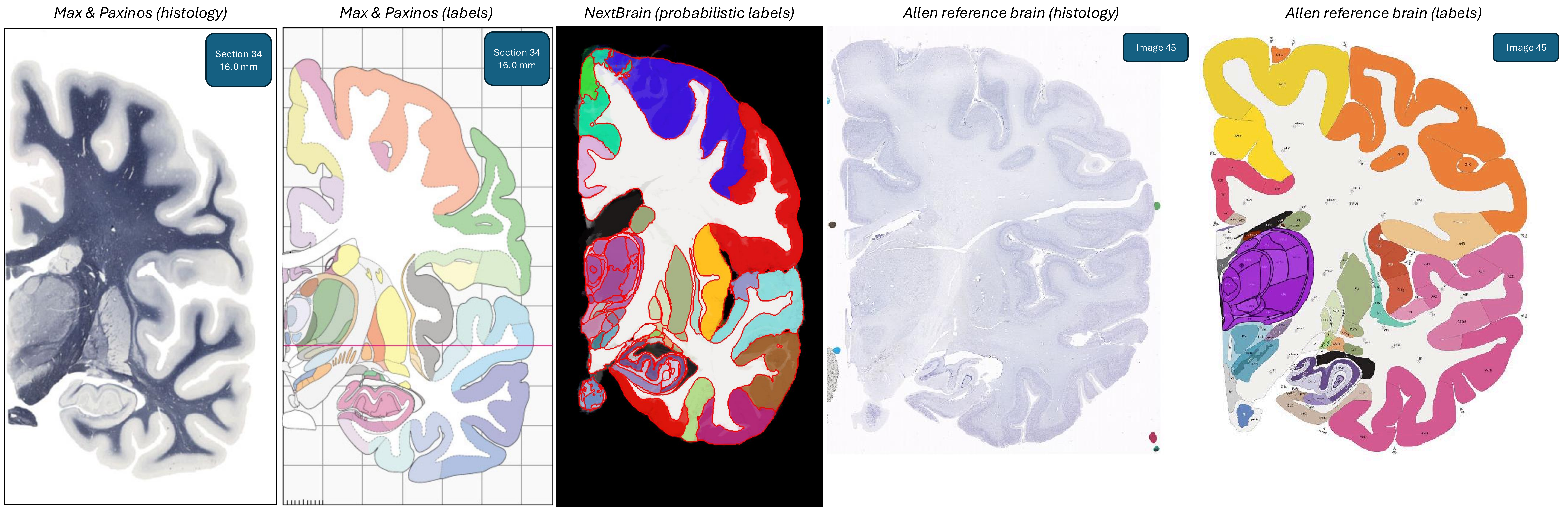
Max & Paxinos (histology)
Section 34
16.0 mm
Max & Paxinos (labels)
Section 34
16.0 mm
NextBrain (probabilistic labels)
Allen reference brain (histology)
Image 45
Allen reference brain (labels)
Image 45

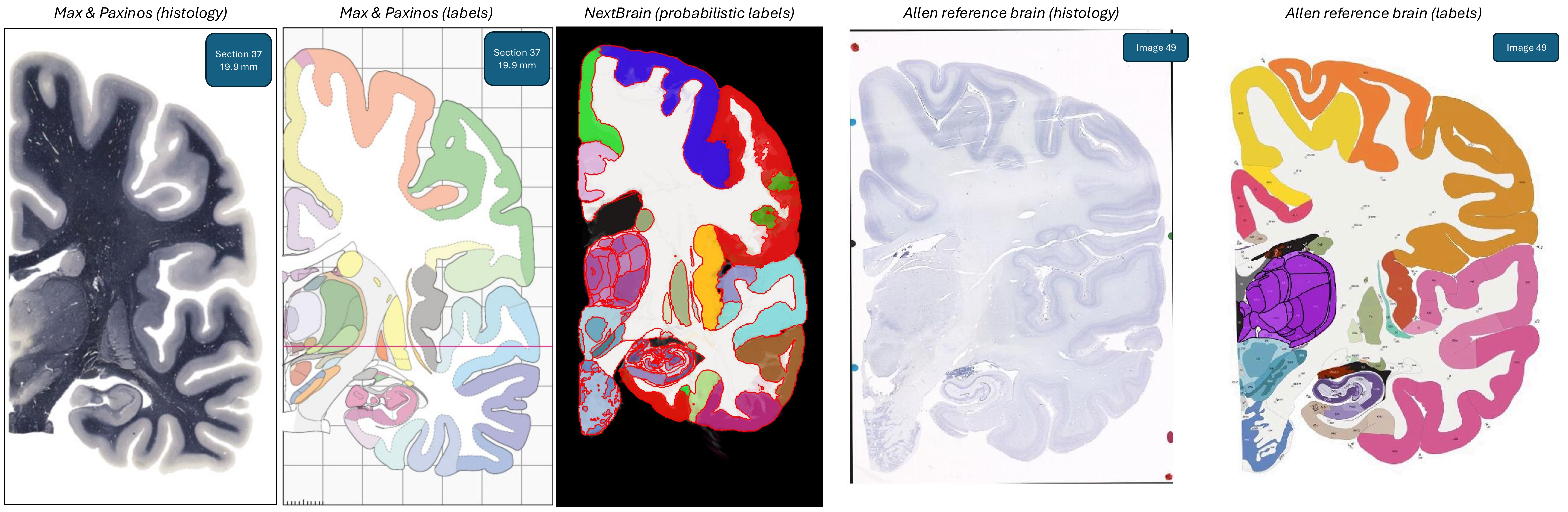
Max & Paxinos (histology)
Section 37
19.9 mm
Max & Paxinos (labels)
Section 37
19.9 mm
NextBrain (probabilistic labels)
Allen reference brain (histology)
Image 49
Allen reference brain (labels)
Image 49

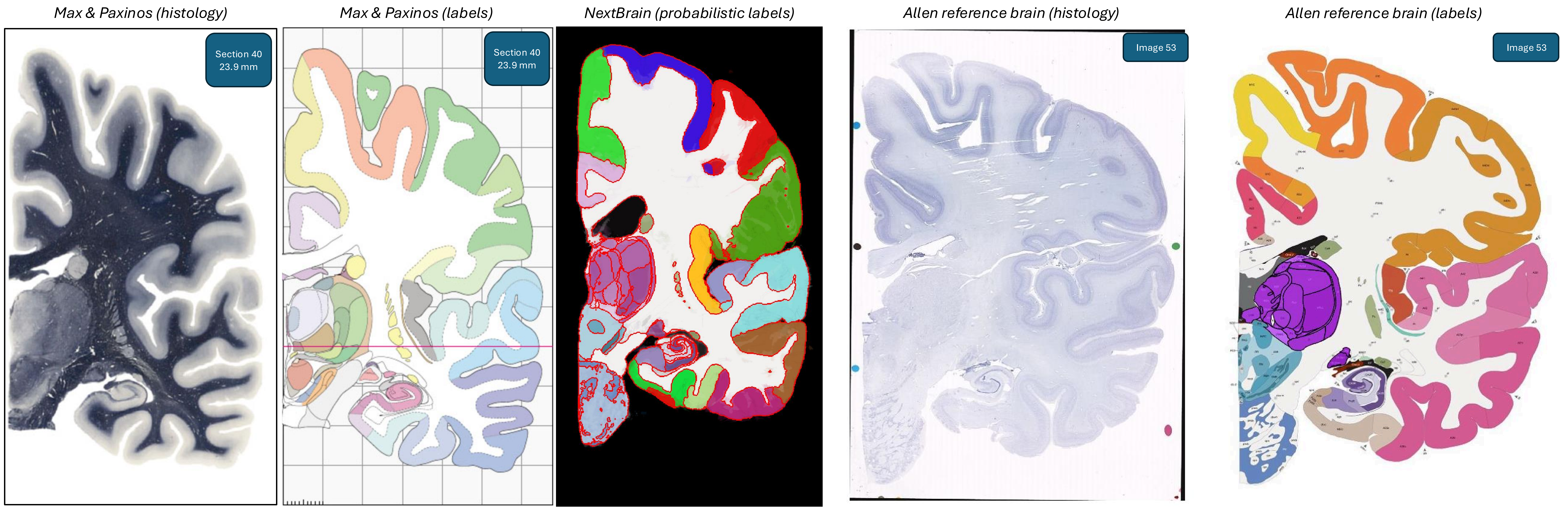
Max & Paxinos (histology)
Section 40
23.9 mm
Max & Paxinos (labels)
Section 40
23.9 mm
NextBrain (probabilistic labels)
Allen reference brain (histology)
Image 53
Allen reference brain (labels)
Image 53

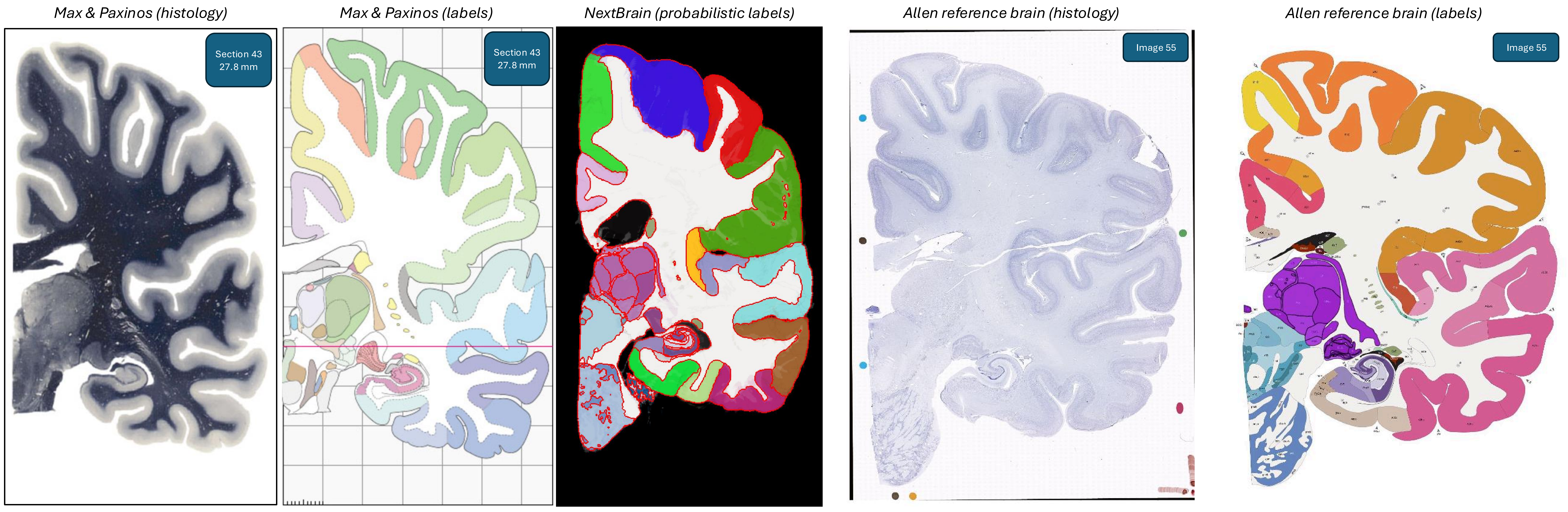
Max & Paxinos (histology)
Section 43
27.8 mm
Max & Paxinos (labels)
Section 43
27.8 mm
NextBrain (probabilistic labels)
Allen reference brain (histology)
Image 55
Allen reference brain (labels)
Image 55

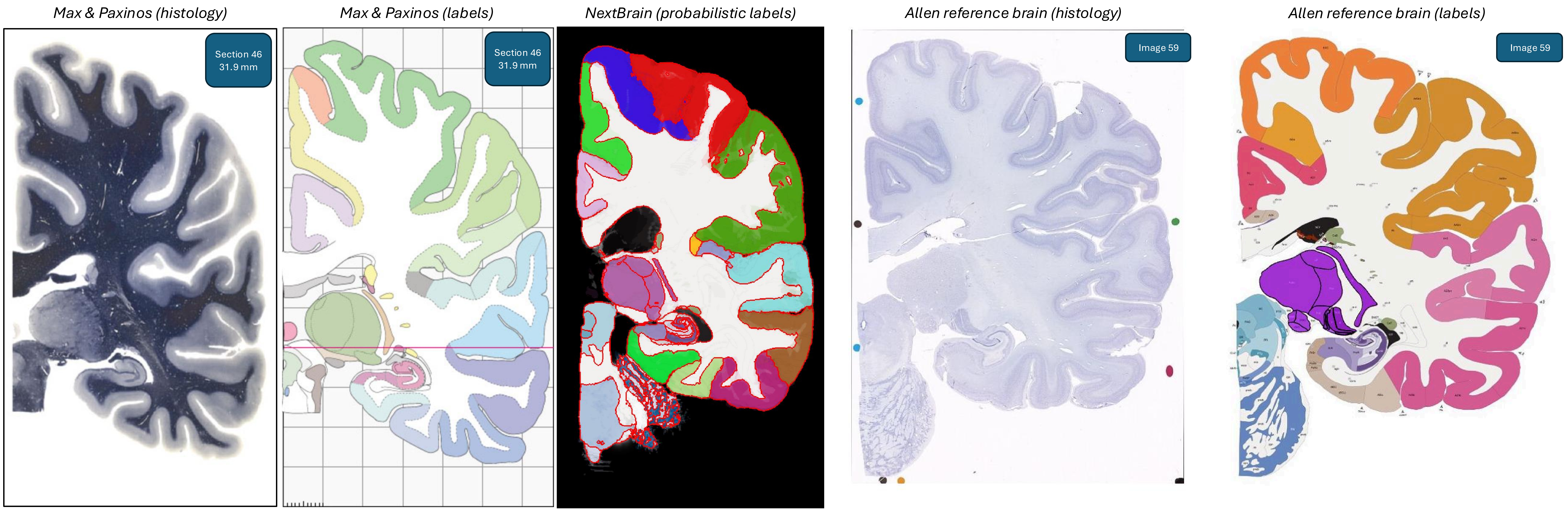
Max & Paxinos (histology)
Section 46
31.9 mm
Max & Paxinos (labels)
Section 46
31.9 mm
NextBrain (probabilistic labels)
Allen reference brain (histology)
Image 59
Allen reference brain (labels)
Image 59

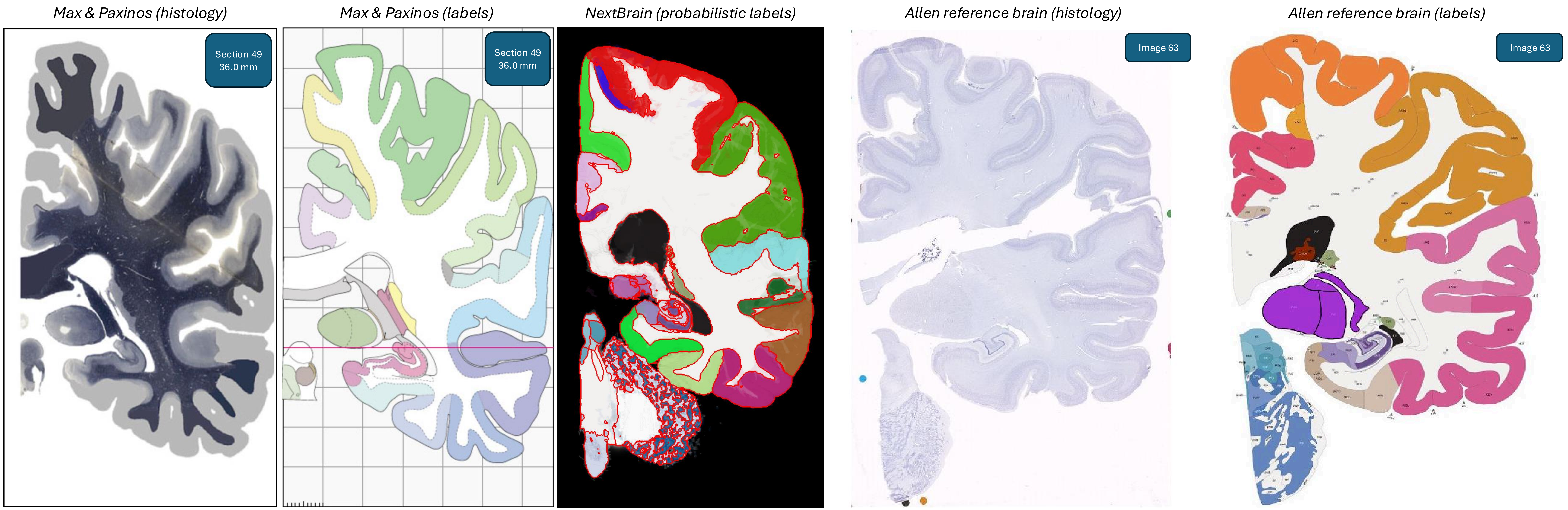
Max & Paxinos (histology)
Section 49
36.0 mm
Max & Paxinos (labels)
Section 49
36.0 mm
NextBrain (probabilistic labels)
Allen reference brain (histology)
Image 63
Allen reference brain (labels)
Image 63

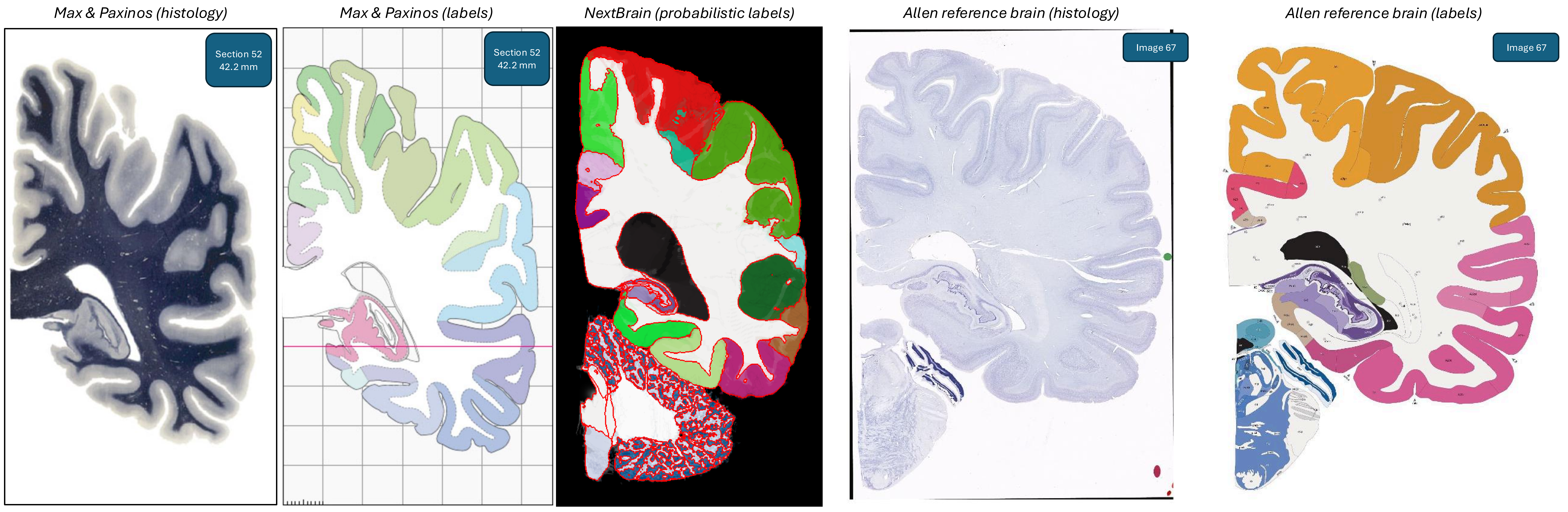
Max & Paxinos (histology)
Section 52
42.2 mm
Max & Paxinos (labels)
Section 52
42.2 mm
NextBrain (probabilistic labels)
Allen reference brain (histology)
Image 67
Allen reference brain (labels)
Image 67

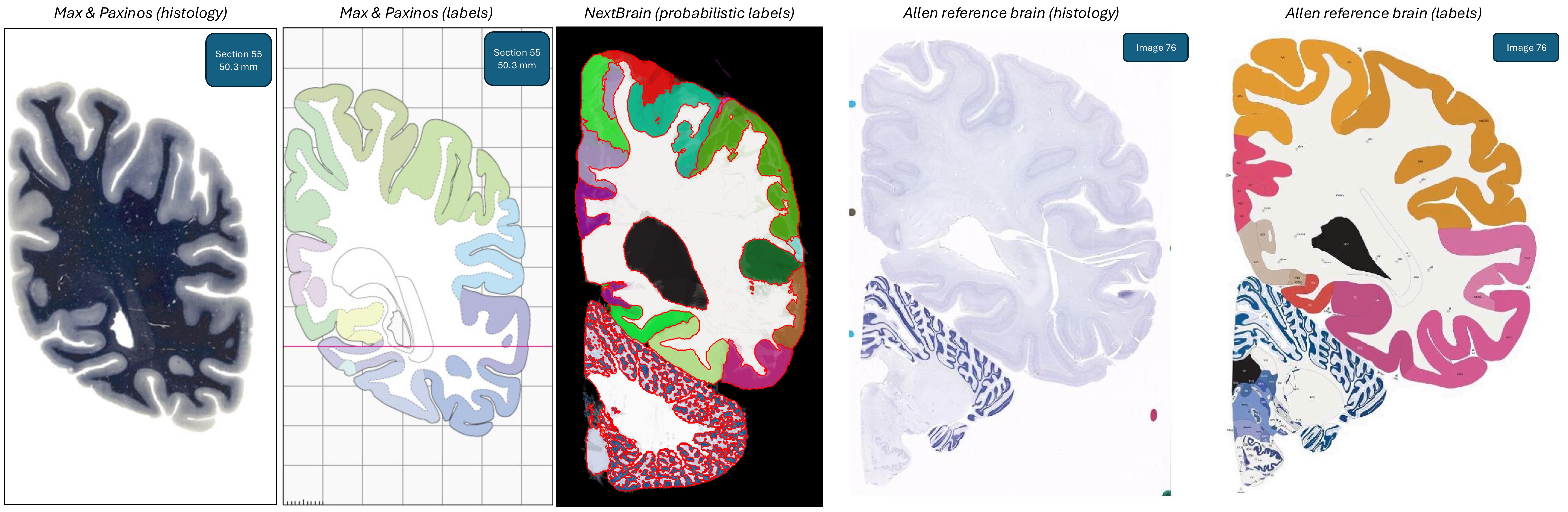
Max & Paxinos (histology)
Max & Paxinos (labels)
NextBrain (probabilistic labels)
Allen reference brain (histology)
Allen reference brain (labels)
Section 55
50.3 mm
Section 55
50.3 mm
Image 76
Image 76

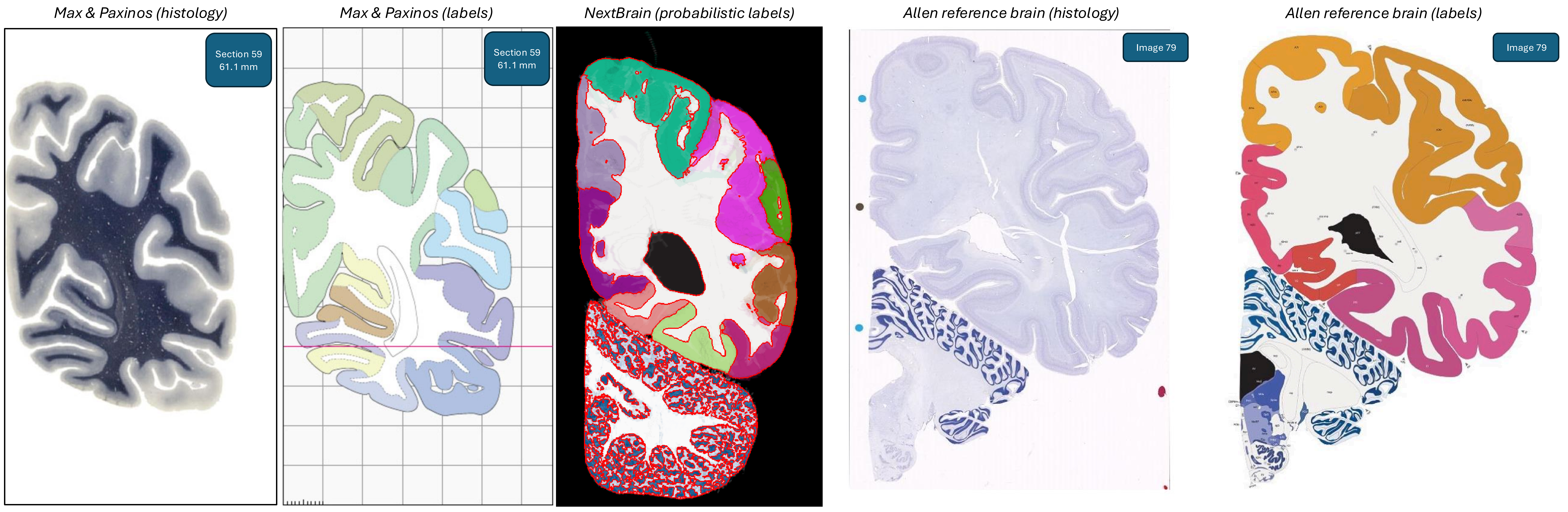
Max & Paxinos (histology)
Section 59
61.1 mm
Max & Paxinos (labels)
Section 59
61.1 mm
NextBrain (probabilistic labels)
Allen reference brain (histology)
Image 79
Allen reference brain (labels)
Image 79

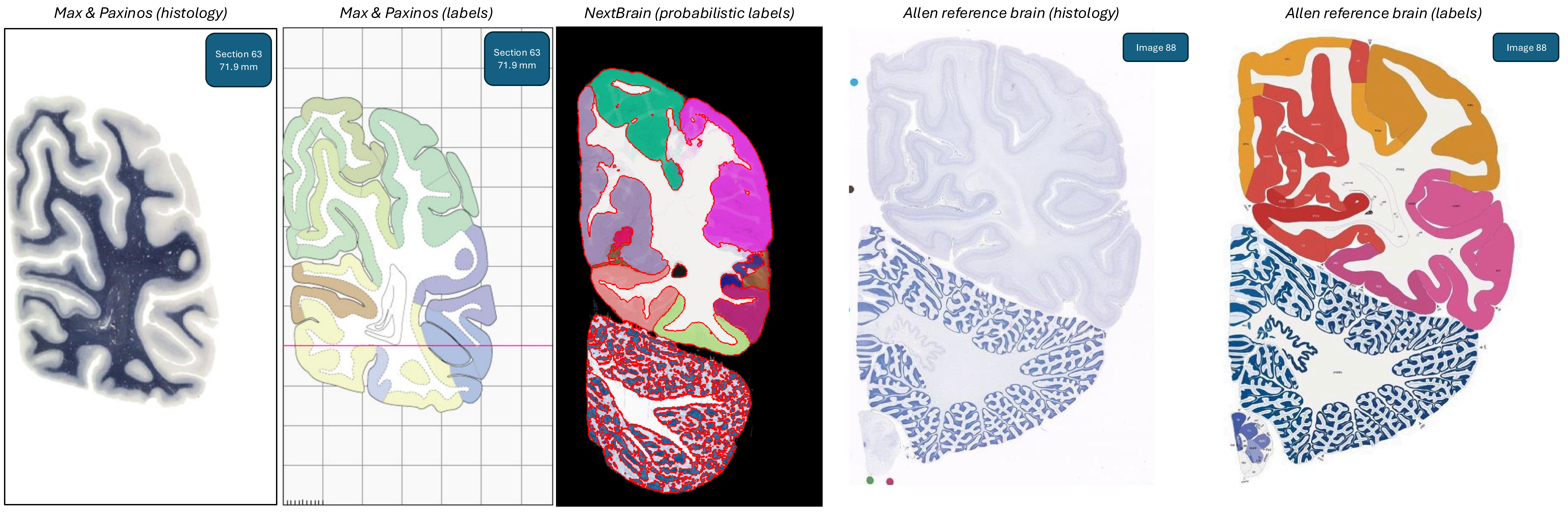
Max & Paxinos (histology)
Section 63
71.9 mm
Max & Paxinos (labels)
Section 63
71.9 mm
NextBrain (probabilistic labels)
Allen reference brain (histology)
Image 88
Allen reference brain (labels)
Image 88

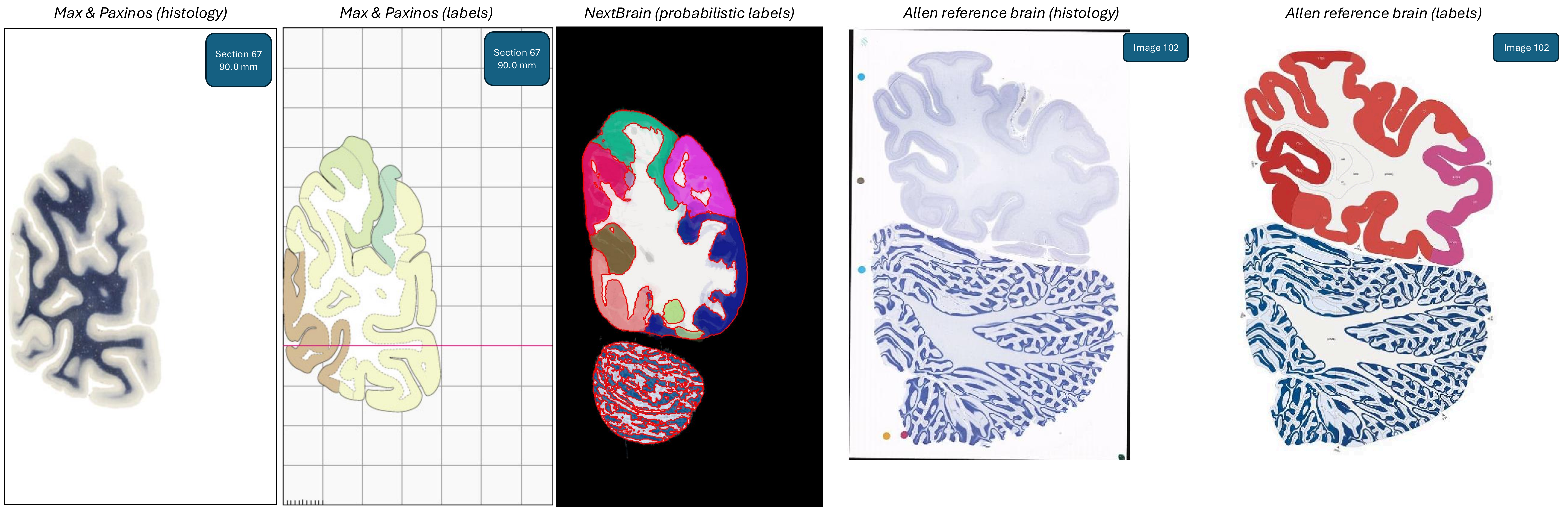
Max & Paxinos (histology)
Section 67
90.0 mm
Max & Paxinos (labels)
Section 67
90.0 mm
NextBrain (probabilistic labels)
Allen reference brain (histology)
Image 102
Allen reference brain (labels)
Image 102
