## Supplementary Methods for "A next-generation, histological atlas of the human brain and its application to automated brain MRI segmentation"

### Manual delineation of brain structures

Our neuroanatomical protocols include 333 regions of interest (ROIs): 34 cortical, and 299 subcortical. The ontology of these ROIs is largely based on the Allen reference brain [1] (henceforth "Allen atlas"). We use their protocol because, leaving aside the sparsity of annotated sections and low sample size (N=1), this atlas leads the pack in terms of number and detail of labels. Our ontology is provided as a spreadsheet in the Supplementary Data. The cortical labels follow the Desikan-Killiany atlas [2]. Our cortical anatomical protocol considers the cerebral cortex as a single ROI in the histology; the subdivision into the 34 ROIs is propagated from the *ex vivo* MRI scans, which were automatically parcellated with FreeSurfer [3]. Our subcortical protocol relies mostly on two atlases: the Allen atlas and the Mai atlas [4]. In addition to these, we used complementary atlases and protocols for specific brain regions, as detailed below; we note that the set of atlases selected to label each region was decided by our senior neuroanatomists (JCA, MB, DK) based on the visibility and clarity of ROIs in our LFB sections.

- *Hippocampus:* In addition to the Allen and Mai atlases, the hippocampal subfields were segmented following [5-8]. The layers of the hippocampus (i.e., stratum lacunosum-moleculare, stratum radiatum, stratum pyramidale, stratum oriens, stratum lucidum) were labelled according to the Allen atlas.

- *Hypothalamus:* The hypothalamus was first segmented into five subregions (anterior superior, anterior inferior, superior tuberal, inferior tuberal, and posterior) as defined in [9,10]. The nuclei within those subregions were then labelled according to [11], as well as the Allen and Mai atlases. Other guiding references were [12-14].

- *Amygdala:* The amygdala was segmented into nine subregions (lateral nucleus, basal nucleus, accessory basal nucleus, central nucleus, medial nucleus, cortical nucleus, anterior amygdaloid area, cortico-amygdaloid transition area), typically visible in *ex vivo* MRI data. The segmentation followed a combination of: *(i)* the Allen and Mai atlases; and *(ii)* a protocol we previously developed for the amygdala atlas in FreeSurfer [15], based on human histology and morphometry resources [16-19].

- *Basal ganglia:* The basal ganglia structures (caudate, putamen and the external and internal segments of the globus pallidus) were traced following the anatomical definitions provided by the Allen and Mai atlases.

- *Basal forebrain:* Structures of the basal forebrain (basal nucleus of Meynert, nucleus of diagonal band, nucleus subputaminalis, substantia innominata) were labelled according to the Allen atlas.

- *Thalamus:* Delineations for 67 thalamic regions were made by three neuroanatomists (M.B., J.A., E.R.), who discussed and agreed to a consistent protocol based on the Allen and Mai atlases, supplemented by [4,20-22] to accommodate for minor heterogeneities in thalamic anatomy – including differences in histological preparation. Following a first round of segmentations, difficult segmentations were reviewed by the more senior neuroanatomist with expertise in thalamic anatomy (M.B.).

- *Midbrain:* The most caudal part of the midbrain was cut with the brainstem, resulting in axial slices, while the cranial part was cut with the cerebrum, resulting in coronal slices. Therefore, we had to adopt two different reference segmentation criteria for the midbrain. For the coronal sections, the primary references used for labelling midbrain regions were the Allen and Mai atlases. For the axial sections, Duvernoy's Atlas of the Human Brain Stem and Cerebellum [23] and [24] were used as additional references.

- *Cerebellum:* The delineation of the molecular and granular layers of the cerebellum as well as cerebellar white matter followed the definitions from the Allen atlas and Duvernoy's Atlas of the Human Brain Stem and CerebelluM [23].

- *Brainstem:* The brainstem was labelled by three neuroanatomists (J.A., E.B., E.R.), who discussed and agreed to a consistent protocol based on the Allen atlas and Duvernoy's Atlas of the Human Brain Stem and Cerebellum [23], as well as on other sources [4,24,25], to accommodate for minor heterogeneities brainstem anatomy and differences in histological preparations.

Brain regions were delineated by our junior neuroanatomists E.R., J.A. (brainstem, midbrain, thalamus, cerebellum and E.B. (hippocampus, brainstem), with guidance and feedback from our senior neuroanatomists D.K. (amygdala), Z.K. (hippocampus, hypothalamus), and M.B. (hippocampus, hypothalamus, thalamus). We note that, by having a single senior neuroanatomist in charge of defining the protocol of each brain region, we maximise the consistency in the annotations – since the junior neuroanatomists in charge of delineating a given region were all trained by the same person.
